# Warming and reduced rainfall alter fungal necromass decomposition rates and associated microbial community composition and functioning at a temperate-boreal forest ecotone

**DOI:** 10.1101/2025.02.20.639335

**Authors:** Anahi Cantoran, François Maillard, Raimundo Bermudez, Artur Stefanski, Peter B. Reich, Peter G. Kennedy

**Author notes:** **Corresponding author:** Anahi Cantoran.

## Abstract

Changes in temperature and rainfall regimes will have significant yet potentially contrasting impacts on rates of soil organic matter (SOM) decomposition. To assess how a combined stress treatment of warming and drought impacts the decomposition of fungal necromass—a fast-cycling soil organic matter (SOM) pool—we incubated *Hyaloscypha bicolor* necromass under both ambient and altered conditions (air and soil warming +3.3°C and ∼40% reduced rainfall) at the B4Warmed experiment in Minnesota, USA. We conducted two multi-week incubations, one assessing mass loss and microbial community composition on decaying necromass after 1, 2, 7, and 14 weeks and the second characterizing the substrate utilization capacities of necromass- associated microbial communities after weeks 1 and 7. Warming and reduced rainfall significantly accelerated the initial rate of necromass decay by ∼20%, but overall mass loss was not different between treatments at the end of the 14-week incubation. The accelerated initial rate of decay paralleled shifts in microbial community composition and activity in the altered plots, demonstrating a higher metabolic capability to utilize C and N substrates early in decomposition but a lower capability later in decay. These findings highlight the dynamic, stage-dependent response of fungal necromass decomposition to altered climate regimes, underscoring the importance of considering both temporal dynamics and the functional capacity of microbial communities when assessing the impacts of climate change on soil carbon and nutrient cycling in forest ecosystems.

## 1. INTRODUCTION

Rising temperatures associated with elevated atmospheric greenhouse gas concentrations are altering the functioning of many ecological communities (García et al., 2018; Reich et al., 2015; Traill et al., 2010). This is particularly pertinent in high-latitude ecosystems, where the effects of increased temperatures are occurring most rapidly (Ito et al., 2020; Serreze et al., 2000). A critical process influenced by warming is the decomposition of organic matter, which plays a central role in carbon (C) and nutrient cycling within ecosystems (Conant et al., 2011; Davidson & Janssens, 2006). Studies have shown that higher temperatures can lead to enhance microbial activity, leading to faster breakdown of plant litter and soil organic matter (SOM) (Bond-Lamberty et al., 2004). This, in turn, can contribute to a positive feedback loop of increased carbon dioxide emissions from soil leading to greater warming (Crowther et al., 2016). Given that soils hold twice the amount of C present in the atmosphere and vegetation combined (Lal, 2004), even small changes to SOM decomposition rates may drive significant additional warming.

Along with warmer temperatures, shifting climatic conditions are also projected to make conditions drier in many higher latitude ecosystems (Seneviratne et al., 2006; IPCC, 2013). Greater water limitation can significantly slow organic matter decomposition due to reduced microbial activity (Manzoni et al., 2012; Smith et al., 2009). Specifically, under drier conditions, microbial metabolism is constrained by lower water availability, which limits the diffusion of substrates and enzymes necessary for organic matter breakdown (Allison & Martiny, 2008). Water stress thus reduces microbial growth and activity, resulting in slower decomposition rates and altered nutrient cycling (Schimel et al., 2007). As such, water availability can play an equally critical role in organic matter decomposition dynamics, potentially slowing soil carbon dioxide release and generating a negative feedback loop.

Given the contrasting trends of warming and drying trends on SOM decomposition dynamics, studying scenarios where temperature and water availability are both manipulated in ecologically realistic ways is crucial to predicting the future functioning of higher latitude ecosystems. Previous research in boreal forests demonstrated that experimental warming and reduced soil moisture inhibited microbial activity and soil respiration (frequently used as proxies of organic matter decomposition) compared to ambient conditions, with a stronger effect of warming when soils were most dry (Allison & Treseder, 2008). While those results suggest increased temperature and decreased moisture may additively limit microbial activity, other studies have found that simultaneous changes in both temperature and moisture may counteract one another. For example, working in an Austrian forest in which temperature and rainfall were factorially altered, Schindlbacher et al. (2012) found that warming stimulated soil respiration, while reduced rainfall hindered it. However, respiration in the combined treatment was not significantly different from that in control plots. More recently, Liang et al. (2024), demonstrated that experimental elevated temperatures stimulated soil respiration in forests at the temperate-boreal ecotone-a transition zone between boreal and temperate forests-but this effect was dependent on soil moisture, showing less stimulation when soil moisture levels were experimentally reduced. Surprisingly, there are very few direct measures of organic matter decomposition in field experiments where temperature and moisture are both manipulated to stimulate near-future climates, limiting our understanding of how altered abiotic conditions will influence decomposition rates and associated microbial community composition and function.

There is a growing recognition that microbial necromass contributes a significant portion to the most persistent C pools in soils, making its decomposition a critical area of study in the context of climate change (Xuan et al., 2024). Specifically, the necromass of fungi has represents a sizable fraction of the microbial-derived C in both particulate and mineral-associated SOM fractions (Angst et al., 2021; B. Wang et al., 2021; Angst et al. 2024), which have turnover times on the scale of years to centuries (Lavallee et al., 2020). Further, fungal necromass has also been shown to be an important source of rapidly cycling nitrogen (N) (Zhang et al., 2018), which can limit SOM decomposition rates (Averill & Waring, 2018). Recent studies have indicated that, like other organic matter inputs, fungal necromass decomposition rates are sensitive to soil warming in forested ecosystems (Fernandez et al., 2019; Liu et al., 2024). Using a litter bag approach, which directly measures necromass mass loss, Fernandez et al. (2019) found that experimentally elevated temperatures significantly increased the decomposition rates of fungal necromass in a forested peatland ecosystem, particularly in microenvironments that experienced the greatest change in water content. While those results indicate that fungal necromass decomposition is sensitive to increased temperature, possibly due to interactions with moisture availability, the unique environmental conditions of peatlands (e.g. the absence of soil, low pH and N availability, low oxygen content due to a shallow water table) (Wright et al., 1992) may be unrepresentative of upland systems. Further, peatlands are known to have a strong filtering effect on microbial decomposer communities (Chen et al., 2022), so determining how more broadly-distributed microbes are impacted by altered environmental conditions is needed to better understand how necromass decomposition and microbial community composition may be linked.

The microorganisms associated with decomposing fungal necromass, known as the fungal necrobiome, are co-dominated by bacteria and fungi (Cantoran et al., 2023; Kennedy & Maillard, 2023). The composition of fungal necrobiome has been consistently shown to represent a distinct subset of the larger soil microbial community that changes over time (Beidler et al., 2020; Brabcová et al., 2016; Fernandez & Kennedy, 2018) and is sensitive to temperature stress. For example, warming has been shown to alter the composition of the fungal necrobiome to favor fast growing bacteria and fungi (Maillard et al., 2022; Zhou et al., 2024). While the composition of the fungal necrobiome is increasingly well characterized, its functioning remains less frequently assessed. Fungal necromass itself is a complex mixture of proteins, polysaccharides, and phenols (See et al., 2021), requiring diverse enzymatic capabilities for breakdown. Studies examining the activity of C- and N-acquiring enzymes have shown that fungal necromass can be a “hotspot” for microbial activity (Brabcova et al., 2016), although exactly which enzymes are most expressed appears to vary depending on the stage of decomposition (Maillard et al., 2021). Specifically, C acquisition may be preferentially targeted during the early stages of necromass decomposition, while N acquisition may be more common later (Maillard et al., 2021). Functional shifts also parallel changes in composition of the fungal necrobiome community, with both ectomycorrhizal fungi and oligotrophic bacteria being increasingly abundant at the later stages of decay (Maillard et al., 2021; Maillard et al., 2022). The extent to which the above enzyme activity patterns are impacted by rising temperatures and declining soil moisture is unknown, representing a critical knowledge gap in efforts to predict microbial processes in a changing climate.

In this study, we assessed the dynamics of fungal necromass decomposition at a boreal-temperate forest ecotone in North America, which is currently experiencing significant climate-related changes in forest functioning and composition (Fisichelli et al., 2014; Reich et al., 2022). We measured fungal necromass mass loss over a 14-week period in open-air experimental plots with increased temperature and decreased rainfall relative to plots with ambient temperature and precipitation. We also characterized the structure of microbial communities associated with decomposing fungal necromass in both plot types at 4 time points during the 14-week incubation, as well as the soil microbial communities at the beginning of the incubation. Further, we deployed a second set of fungal necromass samples in the same plots to assess the functional capacity of the fungal necrobiome community using Biolog ‘EcoPlate’ plates at early (1 week) and later (7 weeks) stages of decomposition. We tested the following three hypotheses: 1) Increased temperature and reduced rainfall will speed the decomposition of fungal necromass, resulting in greater initial rates of decay (k) and a lower recalcitrant mass remaining (A); 2) Increased temperature and reduced rainfall will significantly alter the composition of the fungal necrobiome, favoring fast-growing bacteria and fungi both early and late in decomposition, and 3) Increased temperature and reduced rainfall will favor microbial communities with enhanced capacity to utilize simple forms of C early in decomposition and N and more complex forms of C later in decomposition.

## 2. METHODS

### 2.1 Field site and experimental design

This study was conducted at the University of Minnesota Cloquet Forestry Center in Cloquet, Minnesota, USA, in one of two sites for the B4Warmed (Boreal Forest Warming at an Ecotone in Danger) experiment (46°40046′′ N, 92°31012′′ W). This site has a mean annual air temperature of 4.5°C and mean annual precipitation of 807 mm (Rich et al., 2015). Climate projections for this region over the 21^st^ century predict elevated air temperatures as well as increased annual precipitation, although the summer months will become drier and feature longer periods of extended soil water deficit (Liess et al., 2023). The experiment is composed of 3-meter diameter circular plots, with factorial experimental warming (ambient and +3.3 °C) heated using infrared lamps and soil heating cables and reduced mid-summer rainfall (ambient and 40% reduction) (see Rich et al., 2015; Stefanski et al., 2020, for further details). For our study, we only utilized two of the four treatments: the ambient temperature and ambient rainfall treatment (hereafter referred to as ambient) and the +3.3°C and reduced rainfall treatment (hereafter referred to as altered). All the plots had an open overstory and contained a mixture of randomly assigned 12 native Minnesota tree species as well targeted non-native woody shrubs ranging in age from 2-4 years old.

### 2.2 Site moisture and temperature measurements

Soil volumetric water content (cm3 H2O/cm^3^ soil) from 0-20 cm of each plot was measured hourly using a Campbell Scientific CS-616 probe inserted into the soil at a 45° angle (Rich et al., 2015). Aboveground temperature was measured at mid-canopy height (i.e., roughly the average for all planted tree seedlings in each plot) with thermocouples suspended in acrylic blocks imitating a leaf surface. Belowground temperature was measured using sealed thermocouples at 10 cm soil depth, each monitored continuously (logged every 1 minute and average to 60 min intervals) (Rich et al., 2015).

### 2.3 Necromass production and field incubation

Fungal necromass from *Hyaloscypha bicolor* (formerly *Meliniomyces bicolor)* was generated by growing mycelium on potato dextrose agarose (PDA) for 3 weeks, after which plugs were taken and transferred to PD broth (pH 5) and incubated on orbital shakers at 150 rpm for 30 days at 25 °C. Mycelium was collected, washed, homogenized, and freeze-dried for 3 days at -50 °C to create the fungal necromass (see Pérez-Pazos et al., 2024, for details). *Hyaloscypha bicolor* was used because it is common mycorrhizal fungus found in high latitude forests (Grelet et al., 2009; Kohout et al., 2018) and the composition of its necrobiome has been well characterized in previous studies (Fernandez et al., 2019; Fernandez & Kennedy, 2018; Pérez-Pazos et al., 2024). Necromass was added to 53-micron nylon mesh bags (R510, ANKOM Technology, NY, USA) and heat sealed, with a total of 60 mg (± 1 mg) of necromass added for 24 bags for the first time point (after 1 week, n = 24) and 100 mg (± 1 mg) of necromass was added for the subsequent time points (n = 72) resulting in a total of 96 necromass bags.

Necromass bags were incubated ∼5 cm deep into the soil in four different clusters of each plot. Four separate bags were adjacently placed in each cluster, for a total of 16 necromass bags incubation in each plot (Figure S1). Necromass bags were deployed on June 28th, 2023, and incubated for a total of 1, 2, 7, and 14 weeks. Harvests entailed the removal of one bag from each quadrant at each time point, each bag representing a replicate, transporting the bags back to the University of Minnesota and storing them at -20°C. The following day, the necromass was scraped with a sterile spatula and moved to a sterile 1.5mL microcentrifuge tube and stored at -20°C. To capture the initial microbial community composition at the different sites, small ‘Whirl-Pak’ sterilized bags were filled with soils the day of deployment from an adjacent spot where bags were deployed for each quadrant in each plot (n = 24), transported back to the University of Minnesota and stored at -80°C until further analyses.

### 2.4 Necrobiome substrate utilization

A separate necromass incubation was deployed on August 16th, 2023, using the same *H. bicolor* necromass. Bags were incubated for 1 and 7 weeks to represent early and late decay phases. Bags were deployed at two blocks in for each of the ambient and altered conditions. Four bags were incubated for each cluster of each plot in blocks D and E (n=16 per plot). Three of those necromass bags were pooled by treatment within each block after 1 and 7 weeks of incubation. A total of 200 mg (± 20 mg) of necromass from the pooled bags were aliquoted and then suspended in sterile distilled water (1/20 mv) in a 15 mL conical tube and vortexed for 2 minutes. The remaining bag from each treatment under each block were used to calculate mass loss (n=3 per treatment).

To quantify microbial community substrate utilization, EcoPlates (Biolog, Hayward, CA, USA) were used for each block and treatment (n=4) from the pooled necromass slurries. The supernatant was diluted to 10^-1^ and 100 uL were suspended into the microplate wells and then incubated at 20°C in the dark. The absorbance was measured with a SpectraMax i3x Multi-Mode Microplate Reader at every 24 hours for 120 hours at a wavelength of 590 nm. The measurements were done in triplicate with three blanks (water) per plate. The OD values were corrected by subtracting the blanks OD from the OD from each substrate well, with negative values converted to zero.

Each substrate was grouped into a substrate type category: Amines, Amino Acids, Carbohydrates, Carboxylic Acids, Phenols, and Polymers, respectively (Supplemental Table 1). The substrate average well color development (SAWCD) was calculated from the final timepoint collected (120 hours) using the following equation: SAWCD = ∑ ODi/N, where ODi is the corrected OD value of the substrates within a substrate category and N is the number of substrates in that respective category (Feigl et al., 2017).

### 2.5 Molecular analyses

Bacterial and fungal communities on necromass as well as in soils collected at the beginning of the incubation were identified using high throughput sequencing (HTS). Total gDNA was isolated from necromass (10 ± 1.5 mg) and soil (25 ± 1.5 mg) using the DNeasy PowerSoil Extraction Kit (QIAGEN). Extractions followed manufacturer’s instructions, with the addition of three glass 2 mm beads and a bead beating step for 30 seconds. 16S (bacterial) and ITS2 (fungal) DNA amplicons were generated from extracted DNA at the University of Minnesota Genomics Center (UMGC, Minneapolis, MN). 16S amplicons were generated by a previously published method optimized by the UMGC (Gohl et al., 2016) using a dual-indexed PCR to amplify the V4 region of the bacterial rDNA locus using KAPA HiFi polymerase. ITS2 amplicons were generated using the same dual-index technique with a fungal primer set targeting the ITS2 region of the fungal rDNA locus (forward primer, FSeq2, sequence: TCGATGAAGAACGCAGCG; reverse primer, RSeq (Heisel et al., 2015) and KAPA HiFi Hotstart plus dNTPs (Roche) for amplification. Positive controls were also amplified; the Microbial Community Standard (ZymoBIOMICS) for bacteria and synthetic ITS mock community for fungi (Palmer et al., 2018). Additionally, negative controls were included with no DNA template added. For PCR, 25 cycles were used for 16S and ITS2 amplicon generation. Samples were sequenced on a full NextSeq run (2 × 300 bp Illumina chemistry), with 16S and ITS2 amplicons include in the same run.

Raw sequence files were initially processed with cutadapt (Martin, 2011) to identify all sequence reads with primers and sequencing adapters and remove both. The remaining sequences were filtered, denoised, and merged using the DADA2 algorithm (Callahan et al., 2016) using recommended parameters for 16S (maxN = 0, maxEE = c(2, 4), truncQ = 2) and ITS (maxN = 0, maxEE = c(2, 2), truncQ = 2, minLen = 50). Amplicon sequence variants were assigned taxonomy using the SILVA (v138.1, Quast et al., 2012) and UNITE (v9, Abarenkov et al., 2024) databases for 16S and ITS ASVs, respectively. All forward and reverse .fastq files were deposited in the NCBI Short Read Archive under BioProject ID PRJNAXXXXX (released upon article acceptance).

For each ASV table, sequence reads present in the negative controls was subtracted from the read abundances present in the necromass samples. The mock communities were used to determine to what level sequence reads were found. To assign bacterial trophic mode lifestyles, we classified ASVs belonging to Betaprotebacteria, Gammaproteobacteria, Alphaproteobacteria as copiotrophic, and Acidobacteria, Actinobacteria, and Deltaprotebacteria as oligotrophic based on Trivedi et al. (2018). For fungi, we classified ASVs belonging to the Eurotiales, Hypocreales, Morteriellales, Mucorales, Saccharomycetales, Tremellales and Sporidiales as molds and yeasts based on Sterkenburg et al. (2015) to reflect a r-strategist lifestyle.

Total genomic DNA from the samples used for microbial substrate utilization were also assessed with quantitative real-time PCR (qPCR). The abundance of total bacteria and fungi was assessed using 16S rRNA primers 1401F/968R (Cébron et al., 2008) and 18S rRNA primers FR1/FF390 (Prévost-Bouré et al., 2011), respectively. The qPCR reactions were performed in a 20 uL volume containing 2 ng of DNA, 10 uL of SYBR Green dye, 6.4 uL molecular grade water, and 0.8 uL of each primer at 10 uM. Amplification conditions consisted of 5 min at 95°C, followed by 40 cycles of 20 s at 95°C, 30 s at the primer-specific annealing temperatures (56°C and 50°C for 16S and 18S rRNA, respectively), and 60 s at 72°C on a StepOne Realtime PCR machine (ThermoFisher Scientific, Waltham, MA, USA). For each sample two independent qPCR reactions were performed for bacteria and fungi. The qPCR values are reported as gene log copies per gram of necromass.

### 2.6 Statistical analyses

Given the well-established nonlinear nature of fungal necromass decomposition (See et al., 2021), we fitted the proportion of remaining necromass against incubation time (days) in the ambient versus experimental plots (i.e. those with increased temperature and decreased precipitation) using asymptotic and double-exponential decay models. We found that the best fitting model, selected using Akaike’s information criteria (AIC), was asymptotic. To obtain values for k (the initial rate of decay) and A (the stable fraction remaining to decay), asymptotic models were run on every set of samples, with each set representing a group of necromass bags in each quadrat. One-way ANOVAs were then performed on model-derived k and A values, with treatment as explanatory variable.

To assess changes in microbial community composition on fungal necromass for the field experiment, NMDS and PERMANOVA analyses were conducted using the ‘vegan’ package. Specifically, models compared differences in either bacterial or fungal community composition by treatment, incubation time, and their interaction. We also performed the analyses on the microbial communities present in the necromass incubation deployed in mid-August as well as the initial microbial communities present in soil (in this last case, no time term was included).

To test for treatment and time responses on most common bacteria and fungi colonizing necromass, effect sizes were calculated for the top 12 bacterial and fungal genera based on their occupancy (presence/absence divided by the number of samples) and contribution to community level Bray-Curtis dissimilarity (Shade & Stopnisek, 2019). For each bacterial and fungal genus, Cohen’s d effect size was calculated as the mean relative abundance on necromass when incubated in altered plots minus the mean relative abundance when incubated in ambient plots divided by the pooled standard deviation of relative abundance on necromass incubated in either plot type. Further, we grouped samples into early (days 7 and 14) and late (days 49 and 98) stages of decomposition for Cohen’s d calculations.

To test for differences in microbial community substrate utilization in the 7-week incubation, three-way ANOVAs were conducted to test for differences by compound category, treatment, incubation time, and their interactions. Finally, to determine whether bacterial and fungal log copies per gram of necromass differed across treatments and incubation time in the substrate utilization assay, two-way ANOVAs were conducted. Post hoc analyses for ANOVAs were run using Tukey’s HSD test using the ‘stats’ package and compact letter displays were obtained using the ‘agricolae’ package. All statistical analyses and data visualization were conducted in R version 4.2.3 (R Core Team, 2025).

## 3. RESULTS

### 3.1 Treatment effects on climatic conditions and soil microbial communities

Average temperatures in the altered and ambient plots during the duration of the 14-week incubation were 20.0 ± 1.6 (mean ± 1 SD) and 17.6 ± 1.8 (°C), respectively. The average volumetric water content (VWC) in the altered and ambient plots was 0.12 ± 0.04 and 0.17 ± 0.04 cm^3^ H2O/cm^3^ soil, respectively. Weekly averages for both variables varied over the length of the experiment due to seasonal fluctuations in air temperature and precipitation, although the differences between treatments stayed generally consistent (Figure S1). The soil microbial communities were also significantly different between altered and ambient plots for both bacteria and fungi (Bacteria: PERMANOVA p = 0.0015), Fungi: PERMANOVA p = 0.0001, Figure S2). The warmed and reduced rainfall plots were dominated by the bacterial genera *Bradyrhizobium, Candidatus Udaeobacter*, and *Gaiella* and the fungal genera *Podila*, *Solicoccozyma*, and *Trichoderma*. Control plots were dominated by the bacterial genera *Acidothermus*, and *Bradyrhizobium, Candidatus Udaeobacter*, and the fungal genera *Inocybe*, *Penicillium*, and *Trichoderma* (Supplemental Tables 7). Despite these taxonomic shifts, the proportion of copiotrophic bacteria or fast-growing molds and yeasts did not significantly differ across treatments (Bacteria: χ^2^ = 0.140, df = 2, p = 0.932; Fungi: χ^2^ = 0.796, df = 1, p = 0.372) (Figure S2).

### 3.2 Necromass mass loss

The decomposition of fungal necromass occurred rapidly, with nearly 70% of the initial mass being lost in the first two weeks regardless of plot treatment (Figure 1). During this initial decay stage, however, there was a significant difference in rate, with decay being accelerated by approximately 20% in the altered plots compared to the ambient plots (Figure 1b). Mass loss stabilized after 4 weeks in both treatments and was not significantly different between the two plot types during the later stage of decomposition (Figure 1c).

**Figure 1:**
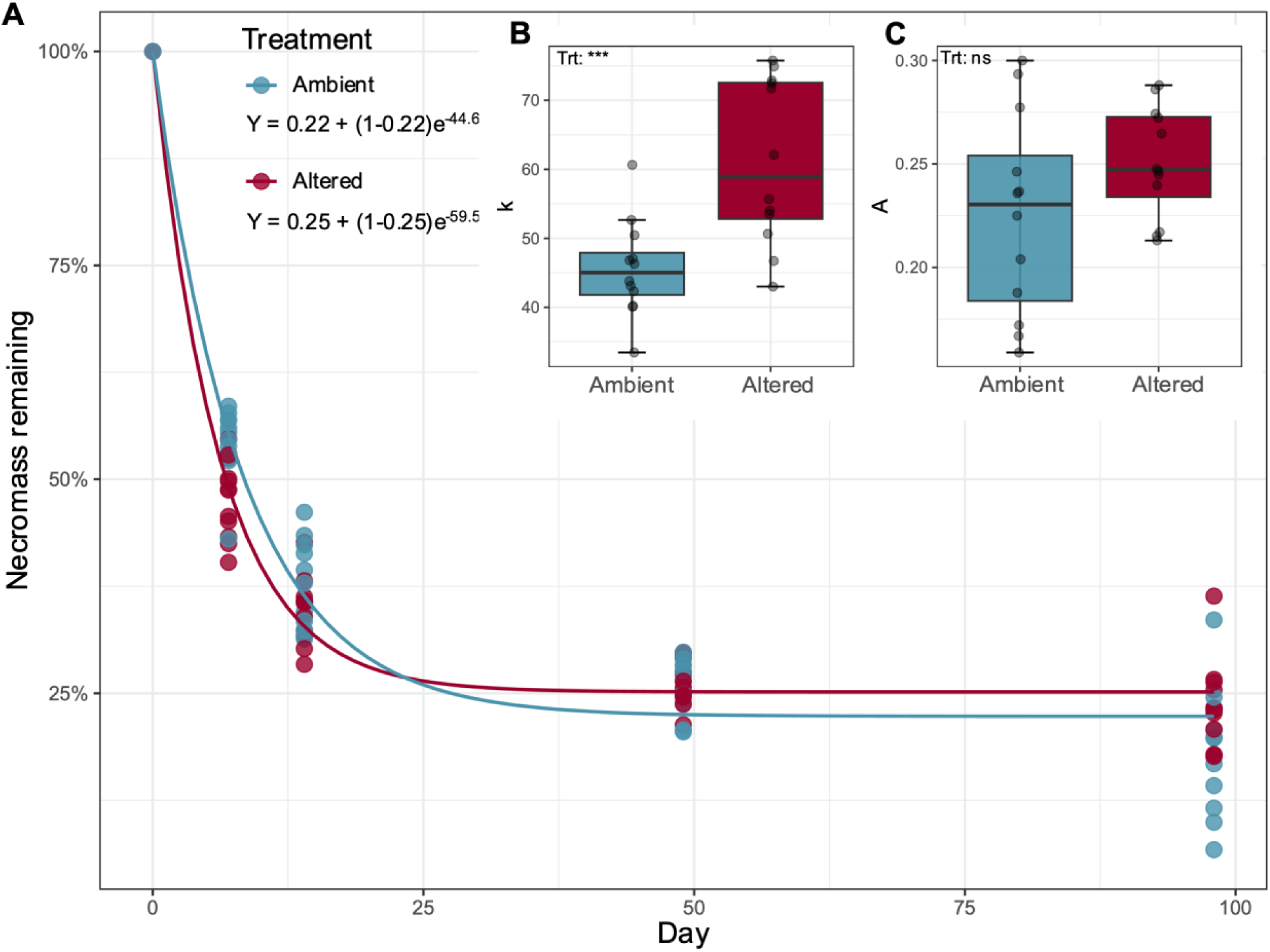
Fungal necromass mass loss during the 14-week field incubation. A) Asymptotic decay model of the percent necromass remaining over time for ambient (blue) and altered (red) conditions. B) Decay rate (k) values and C) Asymptotic parameter (A) values for the ambient and altered conditions. Values were obtained independently for each set of replicates. Statistical significance codes are *p ≤ 0.05, **p ≤ 0.01, ***p ≤ 0.001, and ns for non-significant.

### 3.2 Necrobiome community composition

The composition of the bacterial and fungal communities on decaying fungal necromass differed significantly by treatment and over time (Bacteria: PERMANOVA p = 0.0001, Fungi: PERMANOVA p = 0.0001) (Figure 2a, b). In the altered plots, fungal necromass was highly colonized by bacteria in the genera *Pseudomonas*, *Stenotrophomonas*, and *Chitinophaga*, and fungi in the genera *Fusarium*, *Cunninghamella*, and *Alternaria*, while in the ambient plots necromass was dominated by bacteria in the genera *Paenibacillus*, *Sphingomonas*, and *Mycobacterium* and fungi in the genera *Podila, Exophiala*, and *Humicola*. During early decay (before 14 days), fungal necromass was highly colonized by bacteria in the genera *Pseudomonas*, *Flavobacterium*, and *Stenotrophomonas*, and fungi and the genera *Mucor*, *Fusarium*, and *Humicola*. After 14 days of decomposition, fungal necromass was highly colonized by bacteria in the genera *Mycobacterium*, *Burkholderia*, and *Luteibacter*, and fungi and the genera *Trichoderma*, *Tuber*, and *Podila* (Supplemental Tables 7). The majority of the top 10 most abundant bacterial and fungal genera had similar differences in relative abundance between treatments in early and later stages of decomposition. However, there were a few taxa such as *Pedobacter* and *Trichoderma*, which demonstrated effect sizes in contrasting directions across decay stages (Figure 2c, d). Warming and reduced rainfall increased the proportion of copiotrophic bacteria and fungal molds and yeasts in the necromass communities compared to ambient plots. Notably, this difference in fungal communities was only significant early in the decomposition process (Bacteria Early: χ^2^ = 0.282, df = 2, p = 0.869; Bacteria Late: χ^2^ = 0.191, df = 2, p = 0.909; Fungi Early: χ^2^ = 4.245, df = 1, p = 0.020; Fungi Late: χ^2^ = 2.462, df = 1, p = 0.117) (Figure 2e, f).

**Figure 2:**
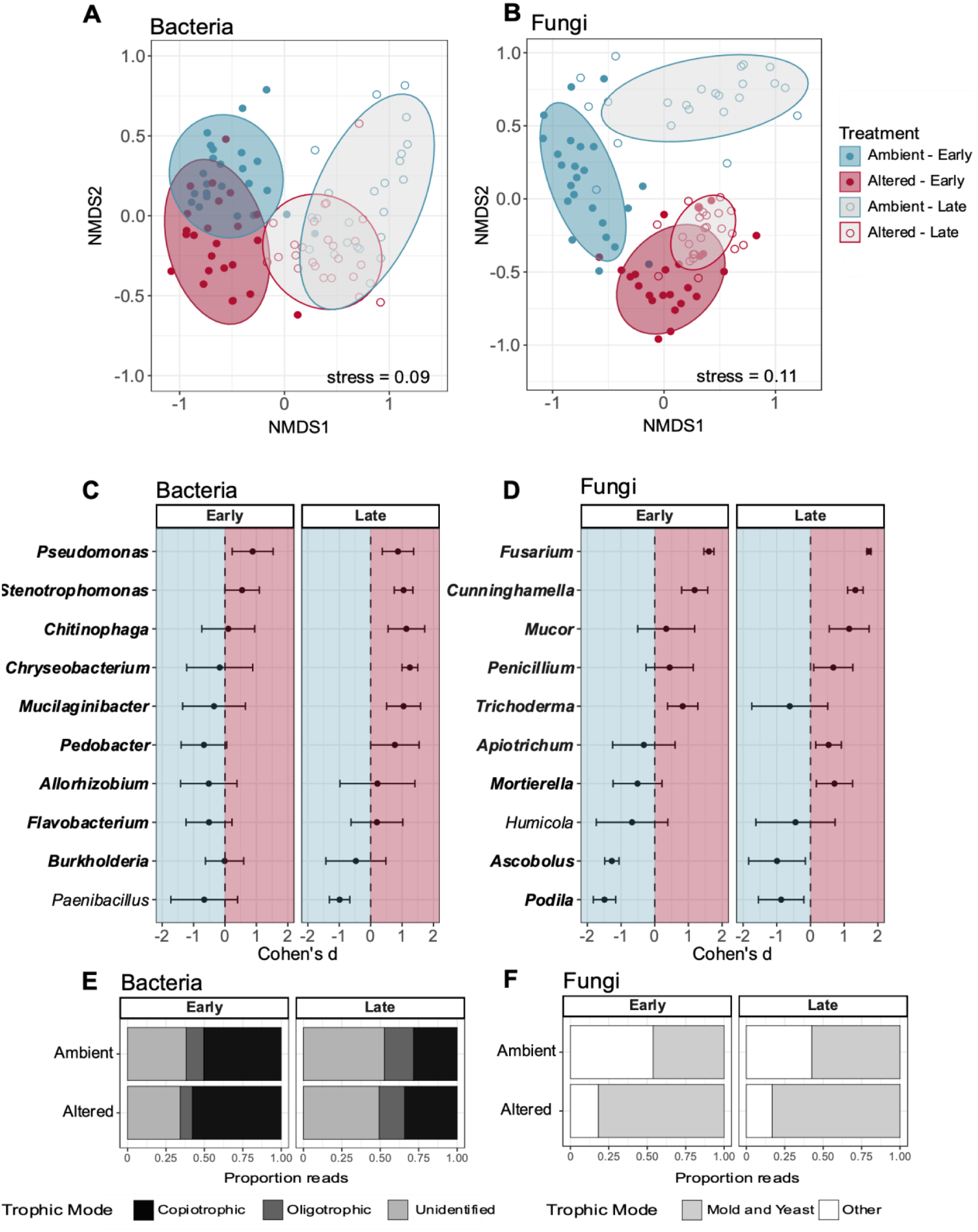
Microbial communities on decaying fungal necromass during the 14-week field incubation. NMDS for A) bacterial and B) fungal communities on necromass. Microbial communities for the ambient conditions are represented in blue and altered conditions are in red. Different time frames are represented as “Early” for necromass incubation at days 7 and 14, and “Late” for necromass incubation at days 49 and 98. Lighter shades of color and open circles represent the late time phase. Cohen’s d values on the C) top 10 dominant bacteria and D) top 10 dominant fungi. Bolded taxa names represent copiotrophic bacteria and fungi classified as molds and yeasts. Proportion of sequence reads of E) bacteria and F) fungi found on necromass classified by their trophic modes across time frames and treatments. Trophic modes for bacteria include copiotrophic (black), oligotrophic (gray), and unidentified (white), while trophic modes for fungi include mold and yeast (black) and other (gray).

### 3.3 Necrobiome substrate utilization

The AWCD values obtained from the substrate utilization incubations were significantly different across substrate (p < 0.0001), time (p < 0.0001), and all interactions (p < 0.01) (Figure 3). AWCD values were higher for microbial communities from the altered plots early in decomposition for 6 of the 7 categories, although only significant for amines and amino acids. Later in decomposition, AWCD values were generally reversed, with all seven categories being higher in the control plots. Amines, amino acids, and phenols all had significantly higher AWCD values for microbial communities from the control plots at this later time point, while no other categories were significantly different. In addition to the general reversal in AWCD values across treatments over time, the values later in decomposition were also consistently lower than early in decomposition, particularly in the altered plots.

**Figure 3:**
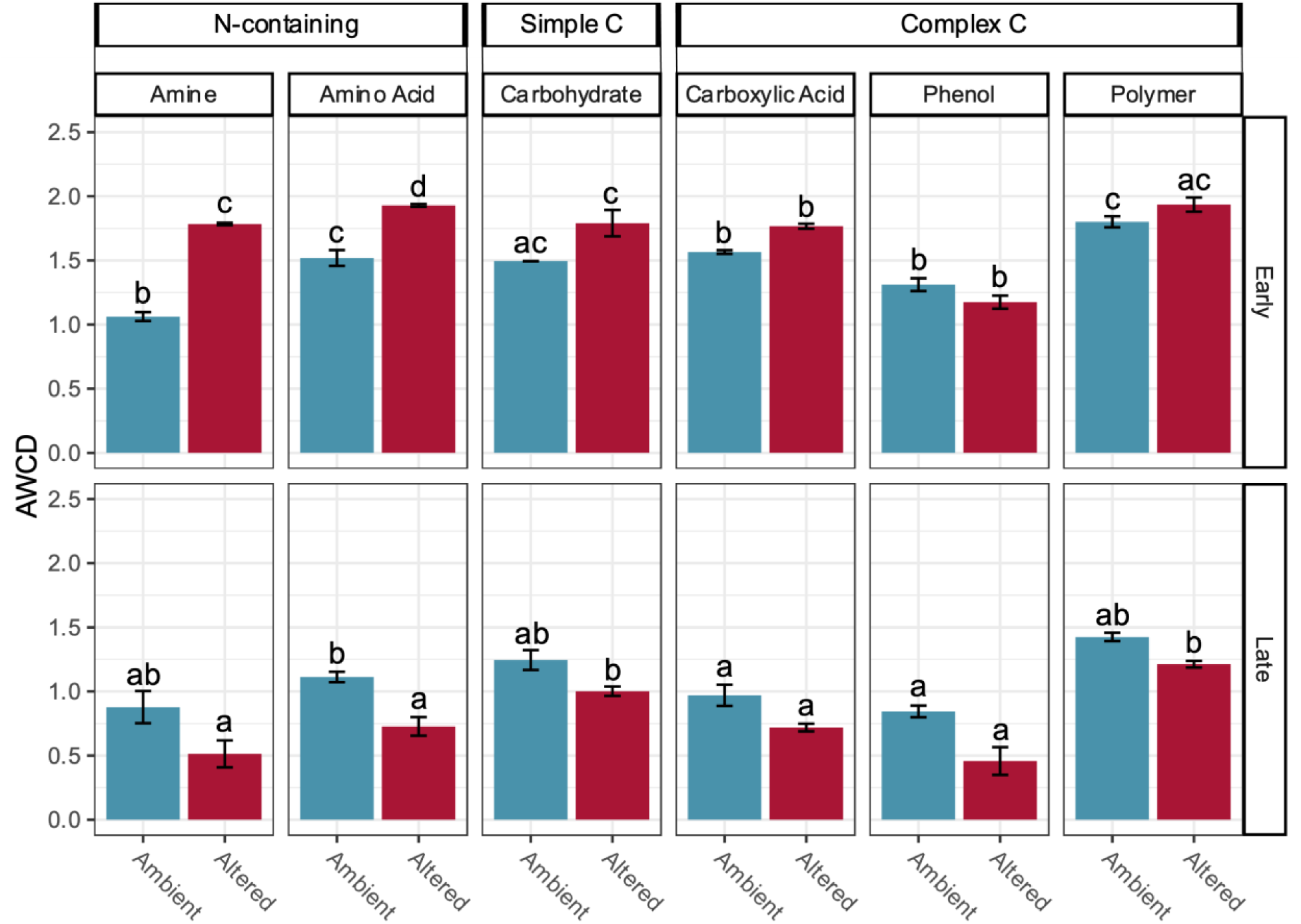
Average well color development (AWCD) from EcoPlates for each substrate type from microbial communities derived from the 7-week incubation. Top row represents the early (day 7) incubation and bottom row is the later (day 49) incubation. Columns from left to right represent different classes of substrates; Amine, Amino Acid, Carbohydrate, Carboxylic Acid, Phenol, and Polymer. See Table S1 for detailed substrate grouping. Significant differences between treatments across the two incubation times from Tukey’s HSD test are indicated with different letters.

During the full 14-week incubation, the microbial communities were significantly influenced by both treatment and time. This was also true for the bacterial and fungal community composition in the 7-week incubation (Figure S3). Though overall microbial communities significantly differed on necromass between the 14-week incubation and the 7-week incubation (Supplemental Tables 3), the top abundant bacteria and fungi were shared across incubations. For example, top abundant bacteria such as *Chitinophaga* and *Pseudomonas,* and fungi such as *Cunninghamella* and *Fusarium* remained within the top 10 most abundant taxa in the 7-week incubation across plot type (Supplemental Tables 7). Additionally, despite differences in composition across time and treatment, the bacterial communities present on the necromass in the 7-week incubation had similar abundances (based on 16S copy number counts) (F1,16 = 0.056, p = 0.815, Figure S4a). In contrast, the fungal communities present on the necromass in the 7-week incubation had significantly higher abundances on necromass in the altered plots compared to ambient plots at both time periods (F1,16 = 6.972, p = 0.0178), Figure S4b). Additionally, the fungal to bacterial (F:B) ratios differed significantly by treatment (F1,16 = 20.144, p = 0.0004) and its interaction with time (F1,16 =12.284, p = 0.003, Figure S4c), demonstrating a shift towards greater fungal-dominated necromass decomposition under warming and reduced rainfall.

## 4. Discussion

Our study reveals a dynamic response of fungal necromass decomposition to warming and reduced rainfall at the temperate-boreal forest ecotone, with distinct patterns emerging across different stages of decay. Specifically, we found that the initial rate of decay was enhanced by ∼20% under warming and reduced rainfall, but that total amount of mass loss was not significantly different between the altered and ambient plots after 14 weeks. This partially supports our first hypothesis of accelerated decomposition, but contrasts with a previous field study in Minnesota, where warming alone enhanced fungal necromass loss in later stages (12-104 weeks; Fernandez et al. 2019). Our later findings do, however, align with a recent global meta-analysis indicating no significant overall impact of either increased temperature or decreased moisture on fungal necromass stocks (Hu et al., 2023), although because we only measured mass loss (a stock output) future research that integrates fungal necromass inputs in this system will be critical for fully understanding the complex dynamics of fungal necromass stocks under climate change.

The stage-dependent response in decomposition likely arises from two key factors. First, because our experiment occurred under field conditions, abiotic factors shifted throughout the study period. As early-stage decomposition coincided with wetter soils (Figure S1), warming likely favored fast-growing microbial taxa that could capitalize on more labile forms of necromass C and climate manipulations did not drive soil moisture to levels where they would overwhelm direct thermal effects (discussed further below). Second, the lack of a significant difference in total mass loss suggests that other factors may counteract the initial acceleration. One likely contributor is the changing chemical composition of the necromass itself. As necromass decomposes, it becomes enriched in recalcitrant compounds (Ryan et al., 2020; See et al., 2021) which are more resistant to microbial breakdown. This increasing recalcitrance, along with greater soil water limitation in the later stage of our experiment, may have slowed decomposition rates, offsetting the initial gains observed under warmer and drier conditions. Our findings highlight the importance of considering microbial community responses and stage-dependent decomposition dynamics when assessing the impacts of climate change on forest ecosystems.

Shifts in soil bacterial and fungal community composition between our experimentally warmed and ambient plots align with previous research in this study system (Fernandez et al., 2023; Van Nuland et al., 2020). While dominant bacterial and fungal genera differed between treatments, overall relative abundance of copiotrophic bacteria, as well as fungal molds and yeasts remained relatively constant in the soil. However, on the necromass itself, we observed notable increases in fungal molds and yeasts in the warmed plots during the early stages of decomposition. Nine of the top 10 dominant fungal genera were classified as molds or yeasts (Figure 3), although some, such as *Ascobolus* and *Podila,* showed significantly greater abundances in the ambient plots. Similarly, nearly all the most abundant bacterial genera were classified as copiotrophic (Figure 2). Given the low C:N ratio of fungal necromass relative to other organic matter inputs (Beidler et al., 2020; DeLancey et al., 2024), it is perhaps unsurprising that fast-growing bacteria and fungi are a dominant part of the necrobiome under conditions highly favorable to growth which supports our second hypothesis.

Consistent with other necromass decomposition studies (Cantoran et al., 2023), we observed significant shifts in the composition of the necrobiome community over time regardless of plot type. Interestingly, the extent of temporal change in the community was greater in the ambient plots for both bacteria and fungi. This could be due to the more rapid initial decomposition of necromass under the combined stress treatment, potentially hastening microbial succession (Tang et al., 2023). Alternatively, the altered plots may have filtered the necrobiome community, limiting compositional changes over time, possibly due to increased network complexity (Yuan et al., 2021). Regardless of the mechanism, these results support our second hypothesis and suggest that warmer and drier climates may favor a necrobiome increasingly dominated by rapidly growing, opportunistic taxa.

Our 7-week incubation provided further insights into the functional capacity of the necrobiome community to utilize a range of C and N sources, including both simple and complex C forms, across early and later decomposition stages. Consistent with the rapid mass loss observed in the 14-week incubation, the week 1 necrobiome community showed significantly higher overall activity than the week 7 community, mirroring patterns observed for C- and N-acquiring enzymes in other studies (Brabcova et al., 2016). Notably, the week 1 necrobiome community in the warmed and reduced rainfall plots exhibited higher activity levels for nearly all substrates compared to the ambient plots. Contrary to our third hypothesis, this difference was most pronounced for N-containing substrates (amino acids and amines), suggesting that fungal necromass may be a particularly important source of labile N. This aligns with previous findings demonstrating rapid N incorporation by bacteria and fungi during necromass decomposition (Maillard et al., 2023) and with observations that most of the N release from fungal necromass occurs within the first 9 days, especially at higher temperatures (X. Wang et al., 2020).

Interestingly, the patterns of C and N utilization reversed at week 7, with the ambient plots showing higher activity levels for all substrates compared to the warmed and reduced rainfall plots. This reversal coincided with significant shifts in microbial community composition, suggesting that later colonizing microbes had a reduced capacity to utilize necromass resources. This may reflect a greater allocation of resources to cellular maintenance under the more stressful conditions of the warmed plots (Romero-Olivares et al., 2019). However, further research, such as culture-based tests of early and later colonizing taxa, is needed to disentangle the relative contributions of intrinsic microbial capacities versus environmental effects. Contrary to expectations based on earlier findings (Brabcova et al., 2016), microbial abundance remained consistent between the two time points, indicating that necromass remains a hotspot of microbial activity even in later decay stages. Furthermore, the higher F:B ratios across treatments and time points that fungi may be more responsive to warming and reduced rainfall than bacteria (Zhou et al., 2024).

Our findings on fungal necromass decomposition have implications for understanding how climate change may alter the dynamics of soil C and N pools. The observed asymptotic decay pattern, with an initial rapid phase followed by slower decomposition, suggests that the processing of fungal necromass could influence the relative size and turnover of these pools. The recalcitrant A fraction, enriched in high molecular weight compounds (Lavallee et al., 2020; Ryan et al., 2020), likely represents a source of more persistent organic matter that could contribute to the particulate organic matter (POM) pool. However, we found no significant difference in the A fraction between our climate treatments, suggesting that later-stage necromass decomposition, and thus its potential contribution to the POM pool, may be relatively insensitive to warming and reduced rainfall. This observation, however, is based on a single necromass type. Since the A fraction size is strongly influenced by cell wall melanin content (Fernandez et al., 2019; See et al., 2021), future research should investigate necromass from species with diverse chemistries to assess the generality of this finding.

In contrast, the significant increase in the rapidly decaying k fraction under warmer and drier conditions could potentially stimulate mineral-associated organic matter (MAOM) formation. This is because the rapid breakdown of the k fraction releases low molecular weight compounds that readily bind to mineral surfaces (Lavallee et al., 2020). However, this potential increase in MAOM formation may be offset by two counteracting factors. First, fast-growing microorganisms, which are favored under warmer conditions, generally have lower carbon use efficiency (CUE) (Domeignoz-Horta et al., 2020). This means they respire a greater proportion of the necromass C, potentially reducing the amount incorporated into the soil. Second, warming can slow MAOM formation (Zhao et al., 2024). Therefore, the net effect of increased necromass input on MAOM stocks remains unclear and directly quantifying the POM and MAOM C and N pools in combination with measures of fungal hyphal production are needed to assess the cumulative effects altered necromass decomposition on soil C and N stocks. Finally, the observed shift towards a more fungal-dominated necrobiome under altered conditions could also potentially influence soil C persistence. Fungi tend to have a higher CUE than bacteria (Soares & Rousk, 2019), and if fungi remain more active under experimental warmed and reduced rainfall conditions (Zhou et al. 2024), more C may be retained in soil (Canarini et al., 2024; Keiblinger et al., 2010) This possibility is supported by other studies demonstrating that soils with high F:B ratios often have higher C stocks (Bailey et al., 2002; Malik et al., 2016).

## Conclusion

Our field-based study demonstrates that fungal necromass decomposition in the boreal-temperate ecotone responds to altered climatic conditions in a dynamic and stage-dependent manner. This ecotone is particularly vulnerable to climate change due to the rapid shifts in temperature and precipitation patterns occurring in these regions (Ito et al., 2020). Our findings show that while warming combined with reduced rainfall initially accelerated decomposition, this effect did not result in significant differences in total mass loss over 14 weeks. By linking these decay dynamics to shifts in microbial community composition and function, our study highlights the critical role of microbial communities in mediating decomposition responses to climate change. Further, by elucidating the links among altered climate regimes, microbial communities, and necromass decomposition dynamics, we emphasize the need for further research to fully assess the consequences of these changes at the boreal-temperate ecotone and similar transitional zones.

## Acknowledgements

We are grateful to many members of the Kennedy lab and B4Warmed team for their assistance with the field and lab experiments. Specifically, we thank Dr. Kaitlyn Beidler for her help with planning, necromass deployment, and manuscript draft revision; Dyonishia Nieves, Soren Mateny, and Hakeem Hamshari for their help with necromass bag deployment and harvesting; Dr. Eduardo Peréz-Pazos for his assistance with necromass generation; Oliv Yesker for her help with harvests and DNA extractions. Funding for this work was provided by a University of Minnesota MnDrive Environment Fellowship to P. Kennedy and A. Cantoran; the National Science Foundation ASCEND Biology Integration Institute (NSF-DBI-2021898), a US Department of Energy, Office of Science, and Office of Biological and Environmental Research award (DE-FG02-07ER64456); Minnesota Agricultural Experiment Station MN-42-030 and MN-42-060; the University of Minnesota College of Food, Agricultural and Natural Resources Sciences and Wilderness Research Foundation, and the Minnesota Invasive Terrestrial Plants and Pests Center through the Minnesota Environment and Natural Resources Trust Fund.

## Supplemental Figures

**Supplemental Figure 1:**
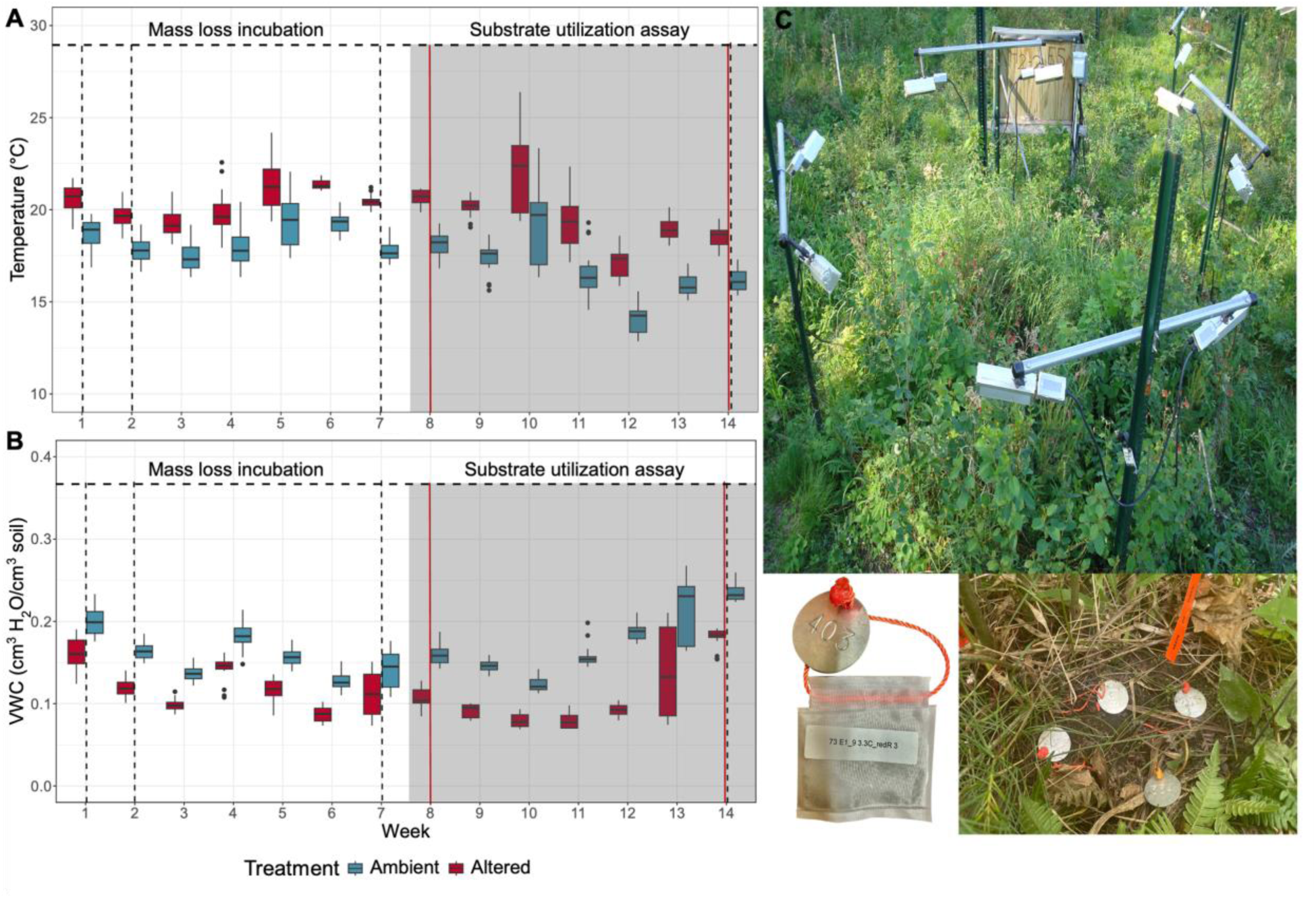
Weekly temperature (A) and volumetric water content (B) averages for the length of the experiment. Black dashed vertical lines represent harvest times of the necromass bags and red solid lines represents harvest times for the necromass bags used for the substrate utilization assays. Week 1 represents one week after initial deployment on June 28^th^, 2023, week 14 represents the last incubation harvest on October 4^th^, 2023, and week 7 (August 16^th^, 2023) was the deployment for necromass bags used for EcoPlates. (C) Images of an altered experimental plot, with a cluster of buried necromass bags and an individual necromass bag pre-incubation.

**Supplemental Figure 2:**
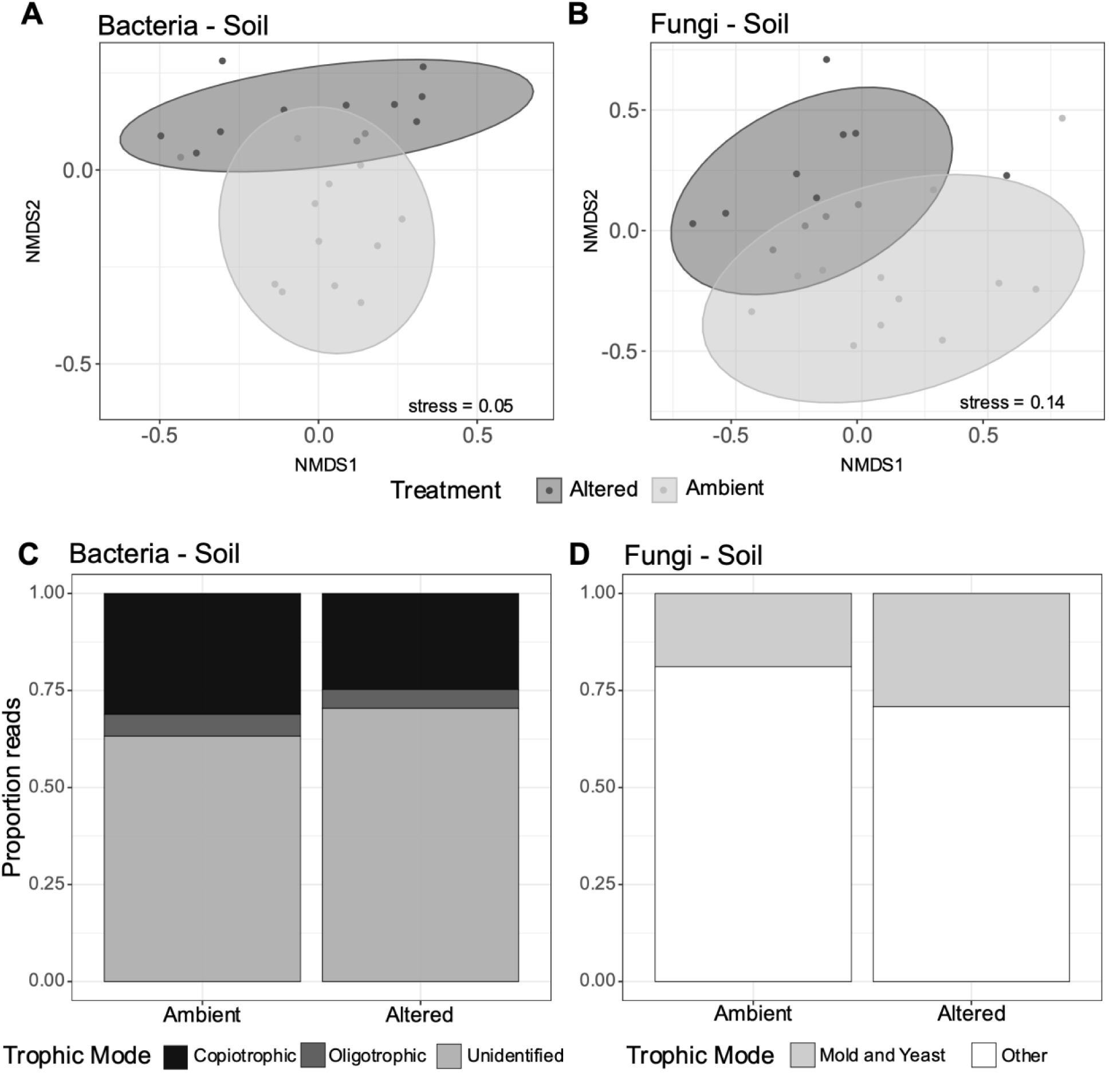
NMDS for A) Bacterial and B) Fungal communities on soil at the initial time of incubation. Proportion of reads of C) bacteria and D) fungi found on soil classified by their trophic modes across treatments. Trophic modes for bacteria include copiotrophic (black), oligotrophic (gray), and unidentified (white), while trophic modes for fungi include mold and yeast (black) and other (gray).

**Supplemental Figure 3:**
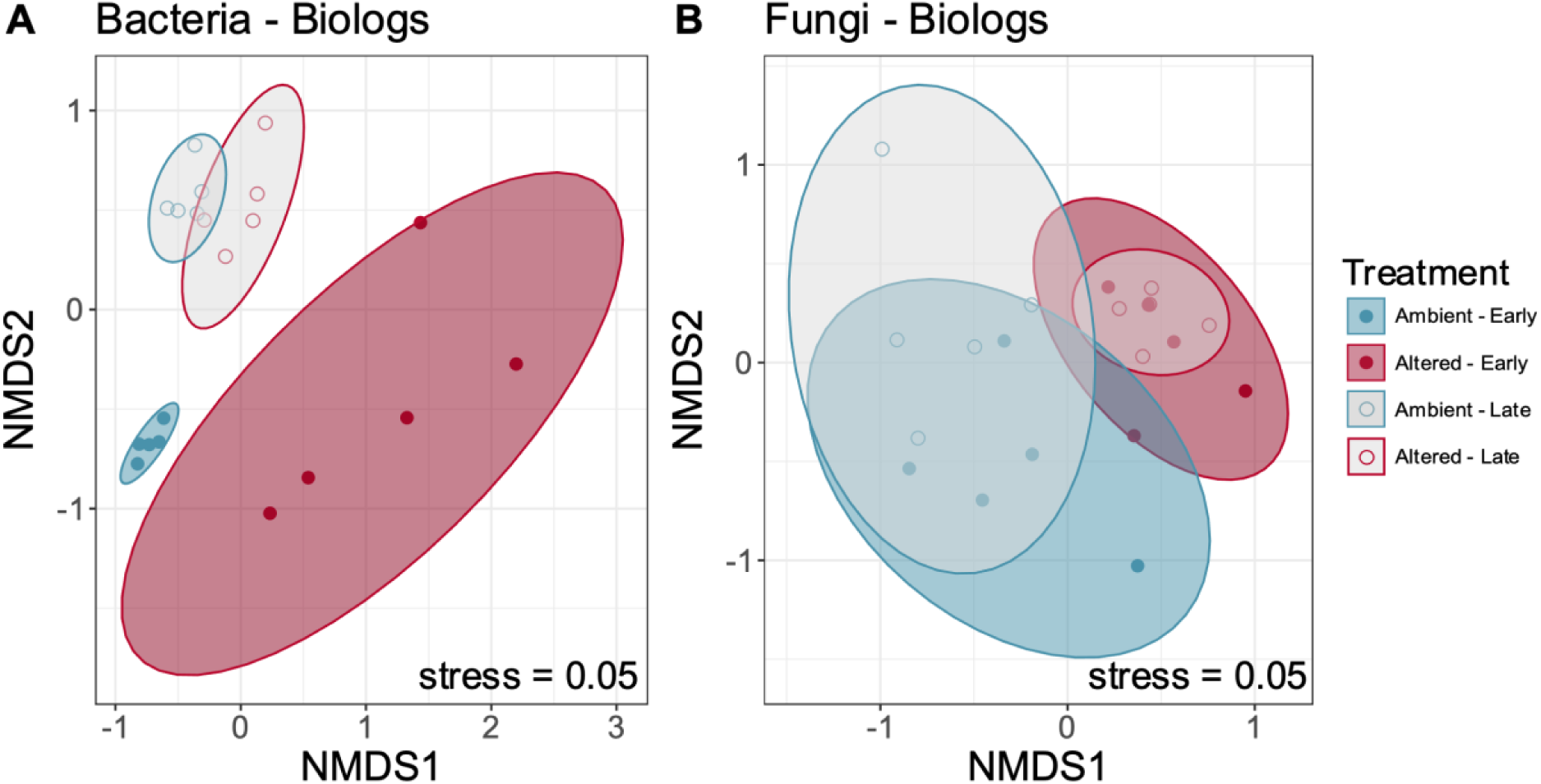
NMDS for A) bacterial and B) fungal communities used in the Biolog EcoPlate Assay. Microbial communities for the ambient conditions are represented in blue and altered conditions in red. Different time frames are represented as “Early” for necromass incubation at day 7 and “Late” for necromass incubation at day 49.

**Supplemental Figure 4:**
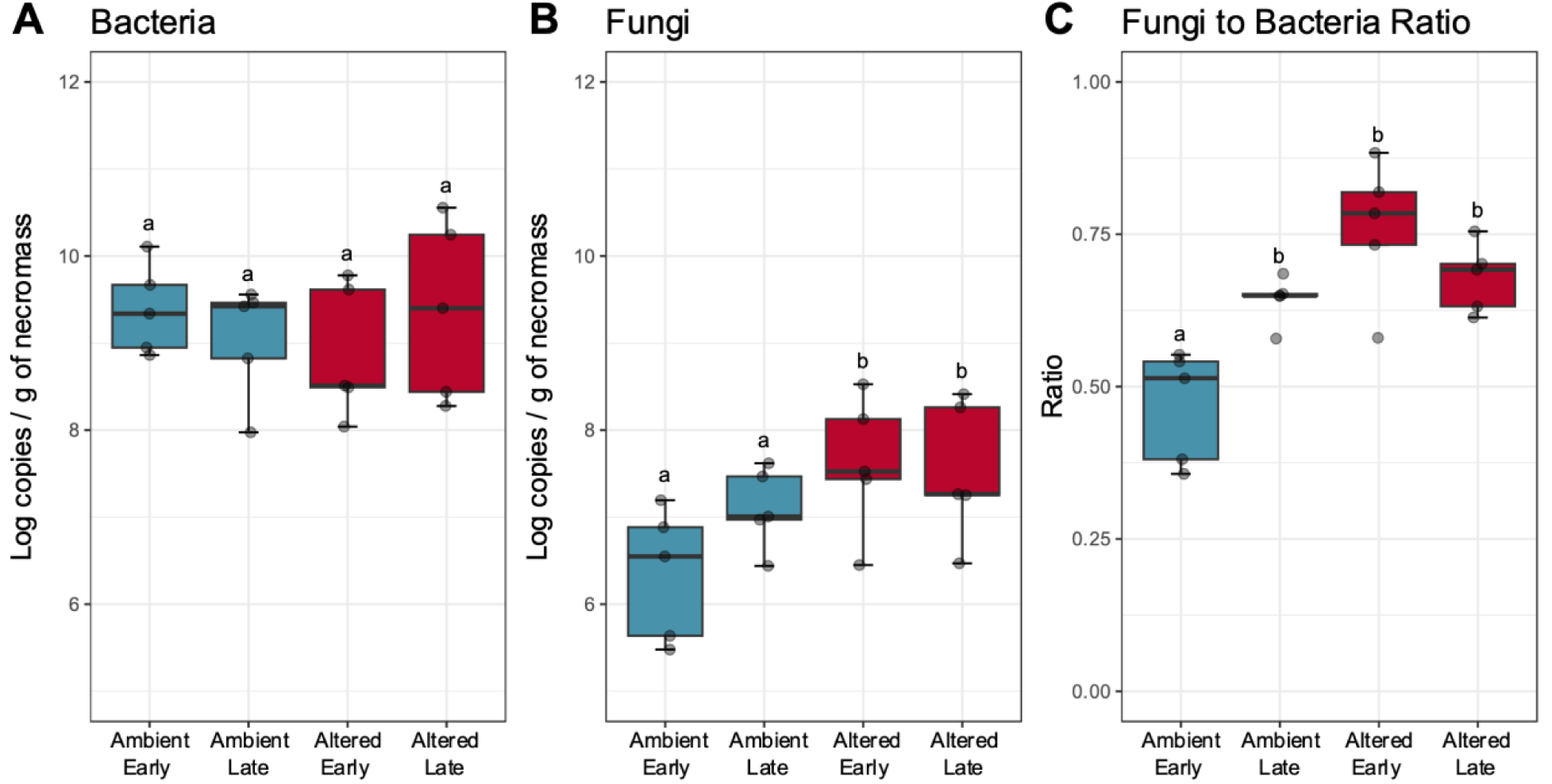
Log number of 16S (bacteria) (A) and ITS (fungi) (B) gene copies obtained by qPCR on fungal necromass. (C) Fungal:bacterial gene copy ratios across treatments. Samples are those used for the substrate utilization assays with the Biolog EcoPlates, which were harvested at day 7 (early) and day 49 (late). Ambient plots are colored in blue and altered plots are colored in red. Significant differences between treatments across the two incubation times from Tukey’s HSD test are indicated with different letters.

## Supplementary Tables

See excel spreadsheet

