## Supplemental Tables for "Warming and reduced rainfall alter fungal necromass decomposition rates and associated microbial community composition and functioning at a temperate-boreal forest ecotone"

### Supplemental Table 1: Classification of substrates in Biolog EcoPlates

| Chemical Group | Substrate | Chemical Class | Chemical Formula |
|---|---|---|---|
| Amines/Amides | Phenylethylamine | N rich | $C_8H_{11}N$ |
| | Putrescine | | $C_4H_{12}N_2$ |
| Amino acids | L-Arginine | N rich | $C_4H_{14}N_4O_2$ |
| | L-Asparagine | | $C_4H_8N_2O_3$ |
| | L-Phenylalanine | | $C_9H_{11}NO_2$ |
| | L-Serine | | $C_3H_7NO_3$ |
| | L-Threonine | | $C_4H_9NO_3$ |
| | Glycyl-L-glutamic acid | | $C_7H_{12}N_2O_5$ |
| Carbohydrates | D-Cellobiose | Simple C | $C_{12}H_{12}O_{11}$ |
| | α-D-Lactose | | $C_{12}H_{12}O_2$ |
| | Methyl-D-glucoside | | $C_7H_{14}O_6$ |
| | D-Xylose | | $C_5H_{10}O_5$ |
| | i-Erythritol | | $C_4H_{10}O_4$ |
| | D-Mannitol | | $C_6H_{14}O_6$ |
| | D,L-a-Glycerol phosphate | | $C_3H_9O_9P$ |
| | Glucose-1-phosphate | | $C_6H_{13}O_9P$ |
| | Pyruvic acid methyl ester | | $C_4H_6O_3$ |
| | N-Acetyl-D-glucosamine | | $C_8H_{15}NO_6$ |
| Carboxylic Acids | D-Galactonic acid lactone | Complex C | $C_6H_{10}O_6$ |
| | D-Galacturonic acid | | $C_6H_{10}O_7$ |
| | γ-Hydroxy butyric acid | | $C_4H_8O_3$ |
| | Itaconic acid | | $C_5H_6O_4$ |
| | α-Keto butyric acid | | $C_4H_6O_3$ |
| | D-Malic acid | | $C_4H_6O_5$ |
| | D-Glucosaminic acid | | $C_6H_{13}NO_6$ |
| Phenols | 2-Hydroxy benzoic acid | Complex C | $C_7H_6O_3$ |
| | 4-Hydroxy benzoic acid | | $C_7H_6O_3$ |
| Polymers | Tween 40 | Complex C | $C_{62}H_{122}O_{26}$ |
| | Tween 80 | | $C_{64}H_{124}O_{26}$ |
| | α-Cyclodextrin | | $C_{36}H_{60}O_{30}$ |
| | Glycogen | | $(C_6H_{10}O_5)n$ |

### Supplemental Table 2: Mass Loss ANOVA

| One-Way ANOVA for treatment effects on k and A | | | | | | |
|---|---|---|---|---|---|---|
| k: initial decomposition rate | | | | | | |
| | Df | Sum Sq | Mean Sq | F value | Pr(>F) | Sig |
| Treatment | 1 | 0.0109 | 0.010897 | 15.2 | 0.000771 | *** |
| Residuals | 22 | 0.01577 | 0.000717 | | | |
| A: Stable fraction | | | | | | |
| | Df | Sum Sq | Mean Sq | F value | Pr(>F) | Sig. |
| Treatment | 1 | 0.00389 | 0.003891 | 2.517 | 0.127 | |
| Residuals | 22 | 0.03401 | 0.001546 | | | |

**Supplemental Tables 3: PERMANOVAS for NMDS community analyses**

| 3a: Bacteria PERMANOVA on field necromass only | | | | | | |
|---|---|---|---|---|---|---|
| | Df | SumOfSqs | R2 | F | Pr(>F) | Sig |
| Treatment | 1 | 1.2396 | 0.06323 | 10.0052 | 0.0001 | *** |
| TimeFrame | 1 | 6.6066 | 0.337 | 53.3217 | 0.0001 | *** |
| Treatment:Ti | 1 | 0.483 | 0.02464 | 3.8982 | 0.0048 | ** |
| Residual | 91 | 11.2749 | 0.57513 | | | |
| Total | 94 | 19.6041 | 1 | | | |

| 3b: Fungi PERMANOVA on field necromass only | | | | | | |
|---|---|---|---|---|---|---|
| | Df | SumOfSqs | R2 | F | Pr(>F) | Sig |
| Treatment | 1 | 4.706 | 0.22796 | 34.4041 | 1.00E-04 | *** |
| TimeFrame | 1 | 2.8434 | 0.13774 | 20.7876 | 1.00E-04 | *** |
| Treatment:Ti | 1 | 0.6471 | 0.03135 | 4.7309 | 7.00E-04 | *** |
| Residual | 91 | 12.4475 | 0.60296 | | | |
| Total | 94 | 20.644 | 1 | | | |

| 3c: Bacteria PERMANOVA on Soil Only ; comp ~ treatment | | | | | | |
|---|---|---|---|---|---|---|
| | Df | SumOfSqs | R2 | F | Pr(>F) | Sig |
| Treatment | 1 | 0.2035 | 0.16309 | 4.2872 | 0.0015 | ** |
| Residual | 22 | 1.0443 | 0.83691 | | | |
| Total | 23 | 1.2478 | 1 | | | |

| 3d: Fungi PERMANOVA on Soil Only ; comp ~ Treatment | | | | | | |
|---|---|---|---|---|---|---|
| | Df | SumOfSqs | R2 | F | Pr(>F) | Sig |
| Treatment | 1 | 0.5929 | 0.09476 | 2.3029 | 1.00E-04 | *** |
| Residual | 22 | 5.6639 | 0.90524 | | | |
| Total | 23 | 6.2568 | 1 | | | |

| 3e: Bacteria PERMANOVA on biologs; comp ~ treatment*time | | | | | | |
|---|---|---|---|---|---|---|
| | Df | SumOfSqs | R2 | F | Pr(>F) | Sig |
| Treatment | 1 | 0.9283 | 0.17947 | 6.9967 | 0.0001 | *** |
| TimeFrame | 1 | 1.5509 | 0.29983 | 11.6892 | 0.0001 | *** |
| Treatment:Ti | 1 | 0.5704 | 0.11029 | 4.2995 | 0.0015 | ** |
| Residual | 16 | 2.1228 | 0.41041 | | | |
| Total | 19 | 5.1724 | 1 | | | |

| 3f: Fungi PERMANOVA on biologs; comp ~ treatment * time | | | | | | |
|---|---|---|---|---|---|---|
| | Df | SumOfSqs | R2 | F | Pr(>F) | Sig |
| Treatment | 1 | 1.2032 | 0.3066 | 9.1587 | 0.0001 | *** |
| TimeFrame | 1 | 0.3805 | 0.09697 | 2.8966 | 0.0126 | * |
| Treatment:Ti | 1 | 0.2386 | 0.06081 | 1.8165 | 0.0836 | . |
| Residual | 16 | 2.1019 | 0.53562 | | | |
| Total | 19 | 3.9243 | 1 | | | |

| 3g: Bacteria PERMANOVA on necromass field vs. biolog day7 ; comp ~ deployment * treatment | | | | | | |
|---|---|---|---|---|---|---|
| | Df | SumOfSqs | R2 | F | Pr(>F) | Sig. |
| Deployment | 1 | 0.8735 | 0.1597 | 8.6452 | 1.00E-04 | *** |
| Treatment | 1 | 0.7962 | 0.14556 | 7.8797 | 1.00E-04 | *** |
| Deployment:' | 1 | 0.7688 | 0.14055 | 7.6082 | 1.00E-04 | *** |
| Residual | 30 | 3.0313 | 0.55419 | | | |
| Total | 33 | 5.4698 | 1 | | | |

| 3h: Fungi PERMANOVA on necromass field vs. biolog day7 ; comp ~ deployment * treatment | | | | | | |
|---|---|---|---|---|---|---|
| | Df | SumOfSqs | R2 | F | Pr(>F) | Sig. |
| Deployment | 1 | 0.4714 | 0.07306 | 3.5263 | 0.0062 | ** |
| Treatment | 1 | 1.7204 | 0.26666 | 12.871 | 0.0001 | *** |
| Deployment:' | 1 | 0.2499 | 0.03874 | 1.8698 | 0.0819 | . |
| Residual | 30 | 4.01 | 0.62154 | | | |
| Total | 33 | 6.4518 | 1 | | | |

| 3i: Bacteria PERMANOVA on necromass field vs. biolog day49 ; comp ~ deployment * treatment | | | | | | |
|---|---|---|---|---|---|---|
| | Df | SumOfSqs | R2 | F | Pr(>F) | Sig |
| Deployment | 1 | 0.2756 | 0.06919 | 2.7168 | 0.0037 | ** |
| Treatment | 1 | 0.4193 | 0.10529 | 4.1342 | 0.0002 | *** |
| Deployment:' | 1 | 0.2449 | 0.0615 | 2.415 | 0.0076 | ** |
| Residual | 30 | 3.0428 | 0.76402 | | | |
| Total | 33 | 3.9827 | 1 | | | |

| 3j: Fungi PERMANOVA on necromass field vs. biolog day49 ; comp ~ deployment * treatment | | | | | | |
|---|---|---|---|---|---|---|
| | Df | SumOfSqs | R2 | F | Pr(>F) | Sig. |
| Deployment | 1 | 0.4801 | 0.0745 | 3.7562 | 0.0046 | ** |
| Treatment | 1 | 1.9263 | 0.2989 | 15.0699 | 0.0001 | *** |
| Deployment:' | 1 | 0.2034 | 0.03156 | 1.5912 | 0.1464 | |
| Residual | 30 | 3.8346 | 0.59503 | | | |
| Total | | | | | | |

### Supplemental Tables 4: Chi-Square tests on fungal and bacterial trophic modes

**On fungal necromass**

| **Bacteria** |
|---|
| X-squared = 7.0808, df = 6, p-value = 0.3134 |
| TimeFrame k |
| X-squared = 0.28157, df = 2, p-value = 0.8687 |
| TimeFrame A |
| X-squared = 0.19128, df = 2, p-value = 0.9088 |

| **Fungi** |
|---|
| X-squared = 9.8527, df = 3, p-value = 0.01986 |
| TimeFrame |
| X-squared = 4.4247, df = 1, p-value = 0.03542 |
| TimeFrame A |
| X-squared = 2.4621, df = 1, p-value = 0.1166 |

**On Soil**

| **Bacteria** |
|---|
| X-squared = 0.14084, df = 2, p-value = 0.932 |

| **Fungi** |
|---|
| X-squared = 0.79628, df = 1, p-value = 0.3722 |

**Supplemental Tables 5: ANOVA on SAWCD**

| Three-way ANOVA of SAWCD ~ Substrate * Treatment * Day | | | | | | |
|---|---|---|---|---|---|---|
| | Df | Sum Sq | Mean Sq | F value | Pr(>F) | Sig |
| Substrate | 5 | 2.146 | 0.429 | 56.867 | 1.60E-12 | *** |
| Treatment | 1 | 0.004 | 0.004 | 0.515 | 0.479731 | |
| Day | 1 | 5.372 | 5.372 | 711.884 | < 2e-16 | *** |
| Category:Trea | 5 | 0.203 | 0.041 | 5.376 | 0.001869 | ** |
| Category:Day | 5 | 0.175 | 0.035 | 4.645 | 0.004171 | ** |
| Treatment:Da | 1 | 1.004 | 1.004 | 133.037 | 2.82E-11 | *** |
| Substrate:Tre | 5 | 0.242 | 0.048 | 6.417 | 0.000644 | *** |
| Residuals | 24 | 0.181 | 0.008 | | | |

BY CLASS

| Three-way ANOVA of CAWCD ~ Class * Treatment * Day | | | | | | |
|---|---|---|---|---|---|---|
| | Df | Sum Sq | Mean Sq | F value | Pr(>F) | Sig |
| Class | 2 | 0.2347 | 0.1173 | 22.204 | 9.27E-05 | *** |
| Treatment | 1 | 0.0009 | 0.0009 | 0.161 | 0.695 | |
| Day | 1 | 2.0386 | 2.0386 | 385.732 | 1.72E-10 | *** |
| Class:Treatm | 2 | 0.0149 | 0.0074 | 1.407 | 0.283 | |
| Class:Day | 2 | 0.0107 | 0.0053 | 1.011 | 0.393 | |
| Treatment:Da | 1 | 0.4347 | 0.4347 | 82.255 | 1.02E-06 | *** |
| Class:Treatm | 2 | 0.0374 | 0.0187 | 3.538 | 0.062 | . |
| Residuals | 12 | 0.0634 | 0.0053 | | | |

| Post Hoc | | |
|---|---|---|
| | SAWCD | groups |
| 3.3C_redR:Da | 1.935325 | a |
| 3.3C_redR:Da | 1.9298111 | a |
| amb_amb :Da | 1.8007833 | ab |
| 3.3C_redR:Da | 1.79057 | ab |
| 3.3C_redR:Da | 1.783825 | ab |
| 3.3C_redR:Da | 1.767119 | abc |
| amb_amb :Da | 1.5659976 | bcd |
| amb_amb :Da | 1.5194278 | bcde |
| amb_amb :Da | 1.49449 | bcde |
| amb_amb :Da | 1.4247875 | cdef |
| amb_amb :Da | 1.3115417 | defg |
| amb_amb :Da | 1.2446017 | defg |
| 3.3C_redR:Da | 1.2120167 | defgh |
| 3.3C_redR:Da | 1.174925 | efghi |
| amb_amb :Da | 1.1130917 | fghi |
| amb_amb :Da | 1.0618417 | ghij |
| 3.3C_redR:Da | 1.00168 | ghij |
| amb_amb :Da | 0.9702071 | ghij |
| amb_amb :Da | 0.8778583 | hij |
| amb_amb :Da | 0.8444083 | ijk |
| 3.3C_redR:Da | 0.7276028 | jkl |
| 3.3C_redR:Da | 0.7189619 | jkl |
| 3.3C_redR:Da | 0.5132833 | kl |
| 3.3C_redR:Da | 0.4578667 | l |

Two-way ANOVA for each susbrate

| Amines | | | | | | |
|---|---|---|---|---|---|---|
| | Df | Sum Sq | Mean Sq | F value | Pr(>F) | Sig |
| Treatment_broad | 1 | 0.0639 | 0.0639 | 4.564 | 0.099502 | . |
| Day | 1 | 1.0578 | 1.0578 | 75.585 | 0.000964 | *** |
| Treatment_broad:Day | 1 | 0.5903 | 0.5903 | 42.179 | 0.002899 | ** |
| Residuals | 4 | 0.056 | 0.014 | | | |

| Compact Letter Display for Category: Amines | | | |
|---|---|---|---|
| ambC_ambR:Day 49 | 3.3C_redR:Day 7 | ambC_ambR:Day 7 | 3.3C_redR:Day 49 |
| "ab" | "c" | "a" | "b" |

| Amino Acids | | | | | | |
|---|---|---|---|---|---|---|
| | Df | Sum Sq | Mean Sq | F value | Pr(>F) | Sig |
| Treatment_broad | 1 | 0.0003 | 0.0003 | 0.057 | 0.823124 | |
| Day | 1 | 1.2937 | 1.2937 | 237.732 | 0.000103 | *** |
| Treatment_broad:Day | 1 | 0.3167 | 0.3167 | 58.198 | 0.001585 | ** |
| Residuals | 4 | 0.0218 | 0.0054 | | | |

| Compact Letter Display for Category: Amino Acids | | | |
|---|---|---|---|
| ambC_ambR:Day 49 | 3.3C_redR:Day 7 | ambC_ambR:Day 7 | 3.3C_redR:Day 49 |
| "a" | "b" | "c" | "d" |

| Carbohydrates | | | | | | |
|---|---|---|---|---|---|---|
| | Df | Sum Sq | Mean Sq | F value | Pr(>F) | |
| Treatment_broad | 1 | 0.0014 | 0.0014 | 0.158 | 0.71142 | |
| Day | 1 | 0.5395 | 0.5395 | 60.282 | 0.00148 | ** |
| Treatment_broad:Day | 1 | 0.1453 | 0.1453 | 16.23 | 0.01575 | * |
| Residuals | 4 | 0.0358 | 0.009 | | | |

| Compact Letter Display for Category: Carbohydrates | | | |
|---|---|---|---|
| ambC_ambR:Day 49 | 3.3C_redR:Day 7 | ambC_ambR:Day 7 | 3.3C_redR:Day 49 |
| "ab" | "c" | "ac" | "b" |

| Carboxylic Acids | | | | | | |
|---|---|---|---|---|---|---|
| | Df | Sum Sq | Mean Sq | F value | Pr(>F) | |
| Treatment_broad | 1 | 0.0013 | 0.0013 | 0.304 | 0.61087 | |
| Day | 1 | 1.3513 | 1.3513 | 326.748 | 5.51E-05 | *** |
| Treatment_broad:Day | 1 | 0.1023 | 0.1023 | 24.741 | 0.00763 | ** |
| Residuals | 4 | 0.0165 | 0.0041 | | | |

| Compact Letter Display for Category: Carboxylic Acids | | | |
|---|---|---|---|
| ambC_ambR:Day 49 | 3.3C_redR:Day 7 | ambC_ambR:Day 7 | 3.3C_redR:Day 49 |
| "a" | "b" | "b" | "a" |

| Polymers | | | | | | |
|---|---|---|---|---|---|---|
| | Df | Sum Sq | Mean Sq | F value | Pr(>F) | |
| Treatment_broad | 1 | 0.0031 | 0.0031 | 0.918 | 0.392368 | |
| Day | 1 | 0.6042 | 0.6042 | 181.183 | 0.000176 | *** |
| Treatment_broad:Day | 1 | 0.0603 | 0.0603 | 18.085 | 0.01313 | * |
| Residuals | 4 | 0.0133 | 0.0033 | | | |

| Compact Letter Display for Category: Polymers | | | |
|---|---|---|---|
| ambC_ambR:Day 49 | 3.3C_redR:Day 7 | ambC_ambR:Day 7 | 3.3C_redR:Day 49 |
| "a" | "b" | "b" | "a" |

| Phenols | | | | | | |
|---|---|---|---|---|---|---|
| | Df | Sum Sq | Mean Sq | F value | Pr(>F) | |
| Treatment_broad | 1 | 0.1368 | 0.1368 | 14.522 | 0.018927 | * |
| Day | 1 | 0.7012 | 0.7012 | 74.408 | 0.000993 | *** |
| Treatment_broad:Day | 1 | 0.0312 | 0.0312 | 3.314 | 0.14279 | |
| Residuals | 4 | 0.0377 | 0.0094 | | | |

| Compact Letter Display for Category: Phenols | | | |
|---|---|---|---|
| ambC_ambR:Day 49 | 3.3C_redR:Day 7 | ambC_ambR:Day 7 | 3.3C_redR:Day 49 |
| "ab" | "ac" | "c" | "b" |

**Supplemental Tables 6: ANOVA on qPCR**

| Two-way ANOVA for bacteria log copies ~ Treatment *Day | | | | | | |
|---|---|---|---|---|---|---|
| | Df | Sum Sq | Mean Sq | F value | Pr(>F) | Significance |
| Treatment | 1 | 0.033 | 0.0332 | 0.056 | 0.815 | ns |
| Day | 1 | 0.032 | 0.032 | 0.054 | 0.818 | ns |
| Treatment:Day | 1 | 0.867 | 0.8668 | 1.474 | 0.242 | ns |
| Residuals | 16 | 9.409 | 0.5881 | | | |

| Two-way ANOVA for fungi log copies ~ Treatment *Day | | | | | | |
|---|---|---|---|---|---|---|
| | Df | Sum Sq | Mean Sq | F value | Pr(>F) | Significance |
| Treatment | 1 | 3.594 | 3.594 | 6.972 | 0.0178 | * |
| Day | 1 | 0.565 | 0.565 | 1.095 | 0.3109 | ns |
| Treatment:Day | 1 | 0.866 | 0.866 | 1.68 | 0.2133 | ns |
| Residuals | 16 | 8.248 | 0.515 | | | |

| Pairwise comparisons | diff | lwr | upr | p adj |
|---|---|---|---|---|
| ambC_ambR:Day 49-3.3C_redR:Day 49 | -0.4315942 | -1.73076 | 0.86757161 | 0.7785697 |
| 3.3C_redR:Day 7-3.3C_redR:Day 49 | 0.08019645 | -1.2189693 | 1.37936225 | 0.9979589 |
| ambC_ambR:Day 7-3.3C_redR:Day 49 | -1.1838043 | -2.4829701 | 0.11536151 | 0.0806573 |
| 3.3C_redR:Day 7-ambC_ambR:Day 49 | 0.51179064 | -0.7873752 | 1.81095643 | 0.6787649 |
| ambC_ambR:Day 7-ambC_ambR:Day 49 | -0.7522101 | -2.0513759 | 0.54695569 | 0.3771889 |
| ambC_ambR:Day 7-3.3C_redR:Day 7 | -1.2640007 | -2.5631665 | 0.03516506 | 0.0579581 |

| Two-way ANOVA for Ratio copies ~ Treatment *Day | | | | | | |
|---|---|---|---|---|---|---|
| | Df | Sum Sq | Mean Sq | F value | Pr(>F) | Significance |
| Treatment | 1 | 0.13354 | 0.13354 | 20.144 | 0.000372 | *** |
| Day | 1 | 0.01073 | 0.01073 | 1.619 | 0.221446 | ns |
| Treatment:Day | 1 | 0.08143 | 0.08143 | 12.284 | 0.002933 | ** |
| Residuals | 16 | 0.10606 | 0.00663 | | | |

**Supplemental Tables 7: Top Bacteria and Fungi on Necromass, Biologs, and Soils**

*Top microbes were defined as those that met the 95th percentile of occupancy and ranked within the 10 most abundant
*grey boxes represent top 10

ON NECROMASS (14-week incubation)

| Top Bacteria | 95% Quantile | 0.8947368 |
|---|---|---|
| Genus | occupancy | rel. abundance |
| Pseudomonas | 0.9473684 | 0.128528928 |
| Flavobacterium | 0.9684211 | 0.061198676 |
| Stenotrophomonas | 0.9052632 | 0.056485474 |
| Luteibacter | 0.9789474 | 0.056241605 |
| Burkholderia-Caballeroni | 0.9789474 | 0.052112803 |
| Allorhizobium-Neorhizobi | 1 | 0.043531466 |
| Mucilaginibacter | 1 | 0.041556532 |
| Paenibacillus | 0.9894737 | 0.03903707 |
| Pedobacter | 0.9578947 | 0.032280088 |
| Chitinophaga | 0.9684211 | 0.031661095 |
| Chryseobacterium | 0.8947368 | 0.017856348 |
| Sphingomonas | 0.9789474 | 0.016643874 |
| Microbacterium | 0.9263158 | 0.010766837 |
| Bacillus | 0.9263158 | 0.006326582 |
| Taibaiella | 0.8947368 | 0.005768419 |
| Variovorax | 0.9684211 | 0.005448414 |
| Tardiphaga | 0.8947368 | 0.005349213 |
| Dyadobacter | 0.9368421 | 0.003835836 |
| Cupriavidus | 0.9368421 | 0.003234088 |
| Streptomyces | 0.9263158 | 0.002838956 |
| Rhodopseudomonas | 0.8947368 | 0.001811238 |
| Bosea | 0.8947368 | 0.001793036 |
| Herbiconiux | 0.9052632 | 0.001586128 |

| Top Fungi | 95% Quantile | 0.5789474 |
|---|---|---|
| Genus | occupancy | rel. abundance |
| Trichoderma | 0.8947368 | 0.18257751 |
| Mucor | 1 | 0.17623975 |
| Fusarium | 0.9368421 | 0.14568292 |
| Humicola | 0.9473684 | 0.10366593 |
| Podila | 0.9473684 | 0.06878111 |
| Penicillium | 0.9684211 | 0.02088333 |
| Apiotrichum | 0.8105263 | 0.02057395 |
| Ascobolus | 0.6631579 | 0.01740773 |
| Cunninghamella | 0.8421053 | 0.01539044 |
| Mortierella | 0.9578947 | 0.01512433 |
| Ypsilina | 0.6 | 0.01400871 |
| Sarocladium | 0.7263158 | 0.0084839 |
| Cladosporium | 0.8421053 | 0.00789932 |
| Exophiala | 0.6736842 | 0.00697182 |
| Pyrenochaetopsis | 0.5789474 | 0.00514839 |
| Clonostachys | 0.6 | 0.00442042 |
| Alternaria | 0.7368421 | 0.00349101 |
| Talaromyces | 0.5789474 | 0.00328548 |
| Cortinarius | 0.5789474 | 0.00231413 |
| Neonectria | 0.6210526 | 0.00161869 |
| Ilyonectria | 0.7157895 | 0.00113895 |

ON BIOLOGS (7-week incubation)

| Top Bacteria | 95% Quantile | 0.8 |
|---|---|---|
| Genus | occupancy | rel. abundance |
| Pseudomonas | 1 | 0.074629426 |
| Streptomyces | 0.95 | 0.07283754 |
| Burkholderia-Caballeronia-l | 0.95 | 0.068138354 |
| Bacillus | 1 | 0.058376986 |
| Chitinophaga | 0.8 | 0.047953558 |
| Allorhizobium-Neorhizobiur | 0.85 | 0.041103819 |
| Pedobacter | 1 | 0.03600845 |
| Mucilaginibacter | 0.8 | 0.035756457 |
| Luteibacter | 0.8 | 0.027445636 |
| Chryseobacterium | 0.8 | 0.026778994 |
| Sphingomonas | 0.85 | 0.012715682 |
| Microbacterium | 0.9 | 0.008294409 |
| Paenarthrobacter | 0.85 | 0.006336928 |
| Novosphingobium | 0.8 | 0.004571476 |
| Bosea | 0.8 | 0.004170067 |
| Paenibacillus | 0.95 | 0.00342035 |
| Pseudarthrobacter | 0.8 | 0.001932666 |
| Rhodococcus | 0.85 | 0.000511555 |

| Top Fungi | 95% Quantile | 0.85 |
|---|---|---|
| Genus | occupancy | rel. abundanc |
| Mucor | 1 | 0.20904345 |
| Fusarium | 1 | 0.18375571 |
| Trichoderma | 0.95 | 0.09677564 |
| Humicola | 1 | 0.05744511 |
| Cunninghamella | 0.9 | 0.04095478 |
| Podila | 0.95 | 0.03858864 |
| Cladosporium | 1 | 0.03745991 |
| Penicillium | 1 | 0.02651004 |
| Sarocladium | 0.85 | 0.02581462 |
| Clonostachys | 0.95 | 0.02544057 |
| Mortierella | 1 | 0.02531678 |
| Alternaria | 0.95 | 0.01658065 |
| Absidia | 0.9 | 0.01277943 |
| Ypsilina | 0.85 | 0.00816575 |

*in red are those belonging to the top 10 bacteria and fungi fror
14-week incubation

ON SOIL

| Top Bacteria | 95% Quantile | 1 | CONTINUED | | |
|---|---|---|---|---|---|
| Genus | occupancy | rel. abundance | Genus | occupancy | rel. abundance |
| Candidatus Udaeobacter | 1 | 0.13093074 | Iamia | 1 | 0.005547764 |
| Bradyrhizobium | 1 | 0.041377644 | Gemmata | 1 | 0.0053822 |
| Acidothermus | 1 | 0.039642814 | Phaselicystis | 1 | 0.005268117 |
| Mycobacterium | 1 | 0.033794127 | Nakamurella | 1 | 0.005243813 |
| Gaiella | 1 | 0.033376766 | JGI 0001001- | 1 | 0.004513256 |
| Candidatus Solibacter | 1 | 0.028297697 | HSB OF53-F0 | 1 | 0.004392117 |
| Conexibacter | 1 | 0.027816634 | Terrimonas | 1 | 0.004142883 |
| Haliangium | 1 | 0.026223578 | Microbacteriu | 1 | 0.003757983 |
| Pseudolabrys | 1 | 0.021425577 | Methylocapsa | 1 | 0.003458187 |
| Rhodoplanes | 1 | 0.021381873 | Cohnella | 1 | 0.00343865 |
| Aquisphaera | 1 | 0.020680256 | Ilumatobacte | 1 | 0.003426465 |
| Acidibacter | 1 | 0.020196979 | Zavarzinella | 1 | 0.003415551 |
| Nocardioides | 1 | 0.018971818 | Ramlibacter | 1 | 0.003400007 |
| Bryobacter | 1 | 0.014260468 | Acidicaldus | 1 | 0.003270842 |
| Candidatus Xiphinematol | 1 | 0.013851747 | Edaphobacul | 1 | 0.003165167 |
| Ellin6067 | 1 | 0.013534024 | Pajaroellobac | 1 | 0.003146609 |
| Roseiarcus | 1 | 0.013229489 | Pirellula | 1 | 0.003045321 |
| CL500-29 marine group | 1 | 0.013194801 | Afipia | 1 | 0.002976699 |
| Gemmatimonas | 1 | 0.012320165 | Luedemannel | 1 | 0.002914558 |
| Reyranella | 1 | 0.011941087 | Angustibacte | 1 | 0.002897671 |
| Hyphomicrobium | 1 | 0.011598232 | Phenylobacte | 1 | 0.002844906 |
| RB41 | 1 | 0.011238393 | Solibacillus | 1 | 0.002831543 |
| Sphingomonas | 1 | 0.010918023 | Occallatibact | 1 | 0.002554855 |
| Paenibacillus | 1 | 0.01053143 | Vicinamibact | 1 | 0.002539419 |
| mle1-7 | 1 | 0.009678257 | Acidipila-Silv | 1 | 0.002524598 |
| Solirubrobacter | 1 | 0.008569386 | Xylophilus | 1 | 0.002410803 |
| Oryzihumus | 1 | 0.00842414 | Lapillicoccus | 1 | 0.001896683 |
| Pedomicrobium | 1 | 0.008133124 | Candidatus N | 1 | 0.001889017 |
| Jatrophihabitans | 1 | 0.007386265 | Candidatus K | 1 | 0.001594374 |
| Pir4 lineage | 1 | 0.007198222 | Aetherobacte | 1 | 0.001585209 |
| MND1 | 1 | 0.00712918 | Kitasatospora | 1 | 0.001555083 |
| Pseudonocardia | 1 | 0.007074685 | Bauldia | 1 | 0.001541597 |
| Burkholderia-Caballeroni | 1 | 0.006929057 | Actinoallomu | 1 | 0.001248035 |
| Clostridium sensu stricto | 1 | 0.006895393 | Singulisphae | 1 | 0.001132279 |
| Piscinibacter | 1 | 0.006836451 | Minicystis | 1 | 0.001017787 |
| Flavobacterium | 1 | 0.006731875 | | | |
| Chthoniobacter | 1 | 0.006673735 | | | |
| Cellulomonas | 1 | 0.006492307 | | | |
| GOUTA6 | 1 | 0.006249925 | | | |
| Bacillus | 1 | 0.006192093 | | | |
| ADurb.Bin063-1 | 1 | 0.006067001 | | | |
| Ferruginibacter | 1 | 0.00567877 | | | |
| Puia | 1 | 0.005640779 | | | |
| Streptomyces | 1 | 0.005598049 | | | |

| Top Fungi | 95% Quantile | 0.9166667 |
|---|---|---|
| Genus | occupancy | rel. abundance |
| Inocybe | 0.9166667 | 0.053731784 |
| Podila | 1 | 0.050085964 |
| Penicillium | 1 | 0.044929022 |
| Trichoderma | 1 | 0.033131193 |
| Solicoccozyma | 1 | 0.030809475 |
| Pseudogymnoascus | 1 | 0.022460364 |
| Fusarium | 0.9583333 | 0.020211665 |
| Tomentella | 1 | 0.015662846 |
| Cladosporium | 0.9583333 | 0.014903315 |
| Humicola | 1 | 0.014773532 |
| Scleroderma | 1 | 0.01476108 |
| Myrothecium | 0.9583333 | 0.013804281 |
| Mortierella | 1 | 0.013262657 |
| Tetracladium | 1 | 0.012939557 |
| Linnemannia | 0.9583333 | 0.012084942 |
| Clonostachys | 1 | 0.010797378 |
| Saitozyma | 1 | 0.009398406 |
| Cortinarius | 1 | 0.009090243 |
| Spizellomycetales_gen_Ince | 0.9583333 | 0.00807226 |
| Exophiala | 1 | 0.00732176 |
| Pyrenochaetopsis | 0.9166667 | 0.007155286 |
| Mucor | 1 | 0.006922691 |
| Talaromyces | 1 | 0.006876426 |
| Pleotrichocladium | 1 | 0.006422699 |
| Gymnostellatospora | 1 | 0.005958287 |
| Laccaria | 0.9166667 | 0.005889679 |
| Pseudoechria | 0.9583333 | 0.005746728 |
| Umbelopsis | 1 | 0.0052179 |
| Truncatella | 1 | 0.004952076 |
| Neonectria | 1 | 0.00461179 |
| Spizellomyces | 0.9583333 | 0.0041339 |
| Lasionectriopsis | 1 | 0.003502461 |
| Leucosporidium | 0.9583333 | 0.003263153 |
| Vishniacozyma | 0.9583333 | 0.002965731 |
| Humicolopsis | 1 | 0.002685296 |
| Ilyonectria | 1 | 0.002428458 |
| Keithomyces | 0.9583333 | 0.001588084 |
| Entoloma | 0.9583333 | 0.001384806 |
| Anungitea | 0.9583333 | 0.001044553 |

**Supplemental Tables 7: Top Bacteria and Fungi on Necromass, Biologs, and Soils --- CONTINUED**

ON SOIL AMBIENT

| Top Bacteria | 95% Quantile | 1 |
|---|---|---|
| otu | otu_occ | otu_rel |
| Candidatus Udaeobacter | 1 | 0.09683546 |
| Bradyrhizobium | 1 | 0.040899578 |
| Gaiella | 1 | 0.035677227 |
| Acidothermus | 1 | 0.032818696 |
| Haliangium | 1 | 0.032050418 |
| Candidatus Solibacter | 1 | 0.031452606 |
| Mycobacterium | 1 | 0.029925577 |
| Pseudolabrys | 1 | 0.027432825 |
| Rhodoplanes | 1 | 0.025058175 |
| Conexibacter | 1 | 0.022349685 |
| Aquisphaera | 1 | 0.02024735 |
| Acidibacter | 1 | 0.019260796 |
| CL500-29 marine group | 1 | 0.015804598 |
| Hyphomicrobium | 1 | 0.015573606 |
| Bryobacter | 1 | 0.015518488 |
| Nocardioides | 1 | 0.015454386 |
| Candidatus Xiphinematol | 1 | 0.01452027 |
| Gemmatimonas | 1 | 0.01439677 |
| Reyranella | 1 | 0.014179333 |
| mle1-7 | 1 | 0.014113306 |
| Ellin6067 | 1 | 0.013889645 |
| Roseiarcus | 1 | 0.013092257 |
| Geobacter | 1 | 0.011976376 |
| Pedomicrobium | 1 | 0.011576878 |
| Pir4 lineage | 1 | 0.01024453 |
| Paenibacillus | 1 | 0.010038374 |
| MND1 | 1 | 0.01000706 |
| RB41 | 1 | 0.009866735 |
| Oryzihumus | 1 | 0.009092733 |
| Flavobacterium | 1 | 0.008911679 |
| Burkholderia-Caballeroni | 1 | 0.008595894 |
| Sphingomonas | 1 | 0.007918503 |
| GOUTA6 | 1 | 0.00779705 |
| Clostridium sensu stricto | 1 | 0.007779901 |
| Puia | 1 | 0.007764222 |
| Chthoniobacter | 1 | 0.007405175 |
| ADurb.Bin063-1 | 1 | 0.007125912 |
| Cellulomonas | 1 | 0.007059287 |
| Phaselicystis | 1 | 0.006801384 |
| Gemmata | 1 | 0.006602911 |
| Piscinibacter | 1 | 0.006522593 |
| Solirubrobacter | 1 | 0.006234921 |
| Jatrophihabitans | 1 | 0.006129112 |
| Ferruginibacter | 1 | 0.006035149 |
| Iamia | 1 | 0.005886226 |
| Pseudonocardia | 1 | 0.005746388 |
| Terrimonas | 1 | 0.005417695 |
| Nakamurella | 1 | 0.005225717 |
| Bacillus | 1 | 0.00507201 |
| JGI 0001001-H03 | 1 | 0.004470803 |
| Pirellula | 1 | 0.004313537 |
| Zavarzinella | 1 | 0.003863738 |
| Ilumatobacter | 1 | 0.003825871 |
| Luedemannella | 1 | 0.003572836 |
| Edaphobaculum | 1 | 0.003559941 |
| Ramlibacter | 1 | 0.003476433 |
| Pajaroellobacter | 1 | 0.003476007 |
| Fimbriiglobus | 1 | 0.003461062 |
| HSB OF53-F07 | 1 | 0.003445343 |
| Microbacterium | 1 | 0.00342658 |
| Acidicaldus | 1 | 0.003312948 |
| Cohnella | 1 | 0.003232153 |
| Afipia | 1 | 0.003066113 |
| Angustibacter | 1 | 0.003049737 |
| Phenylobacterium | 1 | 0.003014838 |
| Tundrisphaera | 1 | 0.002865452 |
| Streptomyces | 1 | 0.002780948 |
| Vicinamibacter | 1 | 0.002778043 |
| Methylocapsa | 1 | 0.002724913 |
| Arenimonas | 1 | 0.002601166 |
| Xylophilus | 1 | 0.002505931 |
| Solibacillus | 1 | 0.002432872 |
| Acidipila-Silvibacterium | 1 | 0.002404493 |
| Occallatibacter | 1 | 0.002327578 |
| IS-44 | 1 | 0.002199867 |
| Bauldia | 1 | 0.002154579 |
| Candidatus Nostocoida | 1 | 0.002145843 |
| Parafilimonas | 1 | 0.001953771 |
| Allorhizobium-Neorhizobi | 1 | 0.001741882 |
| Lapillicoccus | 1 | 0.001738422 |
| Aquicella | 1 | 0.001692728 |
| Candidatus Alysiosphaer | 1 | 0.001690377 |
| Rudaea | 1 | 0.001686724 |
| Legionella | 1 | 0.001672942 |
| Noviherbaspirillum | 1 | 0.001600351 |
| Aetherobacter | 1 | 0.001535396 |
| Kitasatospora | 1 | 0.001499019 |
| Granulicella | 1 | 0.001415873 |
| IMCC26207 | 1 | 0.001371779 |
| Candidatus Koribacter | 1 | 0.001302706 |
| possible genus 04 | 1 | 0.001274603 |
| Minicystis | 1 | 0.001214331 |
| Mesorhizobium | 1 | 0.001206311 |
| Singulisphaera | 1 | 0.001193372 |
| Devosia | 1 | 0.001064367 |
| Actinoallomurus | 1 | 0.000891032 |
| Aminobacter | 1 | 0.000885191 |
| Plot4-2H12 | 1 | 0.000516199 |

ON SOIL TREATMENT

| Top Bacteria | 95% Quantile | 1 |
|---|---|---|
| otu | otu_occ | otu_rel |
| Candidatus Udaeobacter | 1 | 0.16502602 |
| Acidothermus | 1 | 0.04646693 |
| Bradyrhizobium | 1 | 0.04185571 |
| Mycobacterium | 1 | 0.03766268 |
| Conexibacter | 1 | 0.03328358 |
| Gaiella | 1 | 0.0310763 |
| Candidatus Solibacter | 1 | 0.02514279 |
| Nocardioides | 1 | 0.02248925 |
| Acidibacter | 1 | 0.02113316 |
| Aquisphaera | 1 | 0.02111316 |
| Haliangium | 1 | 0.02039674 |
| Rhodoplanes | 1 | 0.01770557 |
| Pseudolabrys | 1 | 0.01541833 |
| Sphingomonas | 1 | 0.01391754 |
| Roseiarcus | 1 | 0.01336672 |
| Candidatus Xiphinematol | 1 | 0.01318322 |
| Ellin6067 | 1 | 0.0131784 |
| Bryobacter | 1 | 0.01300245 |
| RB41 | 1 | 0.01261005 |
| Paenibacillus | 1 | 0.01102449 |
| Solirubrobacter | 1 | 0.01090385 |
| CL500-29 marine group | 1 | 0.010585 |
| Gemmatimonas | 1 | 0.01024356 |
| Reyranella | 1 | 0.00970284 |
| Jatrophihabitans | 1 | 0.00864342 |
| Streptomyces | 1 | 0.00841515 |
| Pseudonocardia | 1 | 0.00840298 |
| Oryzihumus | 1 | 0.00775555 |
| Hyphomicrobium | 1 | 0.00762286 |
| Bacillus | 1 | 0.00731218 |
| Piscinibacter | 1 | 0.00715031 |
| Tumebacillus | 1 | 0.00685313 |
| Clostridium sensu stricto | 1 | 0.00601089 |
| Chthoniobacter | 1 | 0.0059423 |
| Cellulomonas | 1 | 0.00592533 |
| Marmoricola | 1 | 0.00555188 |
| HSB OF53-F07 | 1 | 0.00533889 |
| Ferruginibacter | 1 | 0.00532239 |
| Burkholderia-Caballeroni | 1 | 0.00526222 |
| Nakamurella | 1 | 0.00526191 |
| mle1-7 | 1 | 0.00524321 |
| Iamia | 1 | 0.0052093 |
| ADurb.Bin063-1 | 1 | 0.00500809 |
| Lysinibacillus | 1 | 0.00488496 |
| GOUTA6 | 1 | 0.0047028 |
| Pedomicrobium | 1 | 0.00468937 |
| JGI 0001001-H03 | 1 | 0.00455571 |
| Flavobacterium | 1 | 0.00455207 |
| Subgroup 10 | 1 | 0.00445272 |
| MND1 | 1 | 0.0042513 |
| Methylocapsa | 1 | 0.00419146 |
| Pseudarthrobacter | 1 | 0.00418347 |
| Gemmata | 1 | 0.00416149 |
| Pir4 lineage | 1 | 0.00415191 |
| Microbacterium | 1 | 0.00408939 |
| Phaselicystis | 1 | 0.00373485 |
| Cohnella | 1 | 0.00364515 |
| Puia | 1 | 0.00351734 |
| Ramlibacter | 1 | 0.00332358 |
| Solibacillus | 1 | 0.00323021 |
| Acidicaldus | 1 | 0.00322874 |
| Ilumatobacter | 1 | 0.00302706 |
| Nitrospira | 1 | 0.00302689 |
| Zavarzinella | 1 | 0.00296736 |
| Afipia | 1 | 0.00288729 |
| Terrimonas | 1 | 0.00286807 |
| Pajaroellobacter | 1 | 0.00281721 |
| Occallatibacter | 1 | 0.00278213 |
| Edaphobaculum | 1 | 0.00277039 |
| Angustibacter | 1 | 0.00274561 |
| Phenylobacterium | 1 | 0.00267497 |
| Acidipila-Silvibacterium | 1 | 0.0026447 |
| Xylophilus | 1 | 0.00231568 |
| Vicinamibacter | 1 | 0.0023008 |
| Luedemannella | 1 | 0.00225628 |
| Lapillicoccus | 1 | 0.00205494 |
| Candidatus Koribacter | 1 | 0.00188604 |
| Paenisporosarcina | 1 | 0.00180988 |
| Pirellula | 1 | 0.00177711 |
| Pedococcus-Phycicoccus | 1 | 0.00175883 |
| Aetherobacter | 1 | 0.00163502 |
| Candidatus Nostocoida | 1 | 0.00163219 |
| Kitasatospora | 1 | 0.00161115 |
| Actinoallomurus | 1 | 0.00160504 |
| Terrabacter | 1 | 0.00159907 |
| Streptacidiphilus | 1 | 0.00127897 |
| Singulisphaera | 1 | 0.00107119 |
| Rhodopseudomonas | 1 | 0.00099448 |
| Bauldia | 1 | 0.00092862 |
| Minicystis | 1 | 0.00082124 |
| Dactylosporangium | 1 | 0.00080493 |
| Terracidiphilus | 1 | 0.00076653 |
| Streptosporangium | 1 | 0.00044069 |

ON SOIL AMBIENT

| Top fungi | 95% Quantile | 1 |
|---|---|---|
| otu | otu_occ | otu_rel |
| Podila | 1 | 0.07139682 |
| Solicoccozyma | 1 | 0.042453334 |
| Trichoderma | 1 | 0.027751875 |
| Tomentella | 1 | 0.02465676 |
| Penicillium | 1 | 0.02293836 |
| Pseudogymnoascus | 1 | 0.022653135 |
| Humicola | 1 | 0.016486121 |
| Mortierella | 1 | 0.015237493 |
| Clonostachys | 1 | 0.015131618 |
| Cortinarius | 1 | 0.014304248 |
| Pyrenochaetopsis | 1 | 0.012548999 |
| Cladosporium | 1 | 0.012335811 |
| Tetracladium | 1 | 0.010014627 |
| Saitozyma | 1 | 0.009175323 |
| Gymnostellatospora | 1 | 0.007465473 |
| Mucor | 1 | 0.007436557 |
| Neonectria | 1 | 0.006765061 |
| Pleotrichocladium | 1 | 0.006538846 |
| Spizellomycetales_gen_Ince | 1 | 0.006538766 |
| Truncatella | 1 | 0.005964409 |
| Exophiala | 1 | 0.004934107 |
| Leucosporidium | 1 | 0.00421932 |
| Spizellomyces | 1 | 0.004111682 |
| Talaromyces | 1 | 0.004107784 |
| Umbelopsis | 1 | 0.003709959 |
| Humicolopsis | 1 | 0.003627739 |
| Ilyonectria | 1 | 0.002967748 |
| Lasionectriopsis | 1 | 0.002952605 |
| Cryptosporiopsis | 1 | 0.002582381 |
| Laccaria | 1 | 0.002324926 |
| Paraphaeosphaeria | 1 | 0.002195469 |
| Myrothecium | 1 | 0.002147514 |
| Cladophialophora | 1 | 0.001927045 |
| Anungitea | 1 | 0.001554772 |
| Paramyrothecium | 1 | 0.001238034 |
| GS25_ord_Incertae_sedis_g | 1 | 0.000956851 |
| Scleroderma | 1 | 0.000503476 |

ON SOIL TREATMENT

| Top fungi | 95% Quantile | 1 |
|---|---|---|
| otu | otu_occ | otu_rel |
| Clonostachys | 1 | 0.00646314 |
| Collarina | 1 | 0.00336633 |
| Cortinarius | 1 | 0.00387624 |
| Cunninghamella | 1 | 0.01117198 |
| Entoloma | 1 | 0.00133905 |
| Exophiala | 1 | 0.00970941 |
| Fusarium | 1 | 0.03683945 |
| Ganoderma | 1 | 0.00079035 |
| Gymnostellatospor | 1 | 0.0044511 |
| Humicola | 1 | 0.01306094 |
| Humicolopsis | 1 | 0.00174285 |
| Ilyonectria | 1 | 0.00188917 |
| Inocybe | 1 | 0.05970919 |
| Keithomyces | 1 | 0.00200724 |
| Lasionectriopsis | 1 | 0.00405232 |
| Leptodontidium | 1 | 0.0015064 |
| Linnemannia | 1 | 0.02030212 |
| Mortierella | 1 | 0.01128782 |
| Mucor | 1 | 0.00640883 |
| Neonectria | 1 | 0.00245852 |
| Penicillium | 1 | 0.06691968 |
| Pleotrichocladium | 1 | 0.00630655 |
| Podila | 1 | 0.02877511 |
| Pseudoechria | 1 | 0.00316425 |
| Pseudogymnoascu | 1 | 0.02226759 |
| Saitozyma | 1 | 0.00962149 |
| Scleroderma | 1 | 0.02901869 |
| Solicoccozyma | 1 | 0.01916562 |
| Talaromyces | 1 | 0.00964507 |
| Tetracladium | 1 | 0.01586449 |
| Tomentella | 1 | 0.00666893 |
| Trichoderma | 1 | 0.03851051 |
| Truncatella | 1 | 0.00393974 |
| Umbelopsis | 1 | 0.00672584 |
| Vishniacozyma | 1 | 0.0025512 |
